# Move BeTween modAlities (MBTA) employs flow matching to predict single cell data modalities

**DOI:** 10.64898/2026.08.05.743110

**Authors:** Bingxian Xu, Yiyang Zhang, Franziska Michor

## Abstract

Integrating diverse molecular modalities to obtain a comprehensive view of cellular identity remains a major challenge in single-cell biology. A fundamental but underappreciated obstacle is structural mismatch — the phenomenon in which the neighborhood structure of a cell differs depending on which molecular modality is used to define it. Existing approaches typically embed modalities into a shared latent space, which actively erases the structural differences between modalities that make multimodal measurements scientifically valuable. Here we introduce Move BeTween modAlities (MBTA), the first framework explicitly designed to address structural mismatch. Rather than forcing modalities into a shared representation, MBTA maintains modality-specific latent spaces and connects them via flow matching, preserving the structural integrity of each modality while enabling accurate cross-modal translation. Across extensive benchmarks on multi-modal single-cell datasets, MBTA consistently outperformed existing methods, with the largest gains observed in datasets with pronounced structural mismatch. Applied to joint genomic and transcriptomic profiles of breast cancer patients, MBTA identified transcriptomic lineage relationships corroborated by genomic variation and outperformed state-of-the-art transcriptomics-based copy number inference methods. Extending this framework to mouse embryonic development, we reconstructed temporal trajectories jointly defined by gene expression and seven complementary epigenetic modalities. MBTA can connect any number of molecular readouts without erasing their individual character, serving as the computational foundation for assembling multi-layered portraits of cells.

## Introduction

Cells are autonomous machines capable of completing complicated tasks such as dividing and differentiating under the choreographed interplay of different chemical species [1, 2, 3, 4]. Through the advancement of single-cell technologies, the state of single cells can now be resolved through profiling of transcriptomics, proteomics, DNA accessibility and other modalities [5, 6, 7]. In the universe of single-cell profiling modalities, single-cell RNA sequencing (scRNA-seq) remains the most prevalent, with large numbers of cells measured in each scRNA-seq experiment, allowing for the identification of hidden low-dimensional structures [8, 9, 10, 11] such as trees commonly observed in differentiating systems [12, 13, 9]. While scRNA-seq enables precise categorization of cellular states in the transcriptomic space, there are examples in which gene expression alone seems insufficient to fully explain the functional state of a cell; for instance, a recent study showed that neurons with similar gene expression can diverge in both shape and visual response [14]. This observation was echoed in another study demonstrating that drug-induced morphological changes are not reflected in transcriptomic space [15].

To gain a more holistic view of the state of each cell, multi-modal single-cell technologies have started to emerge, offering increasingly sophisticated ways to characterize cell states through the co-profiling of multiple modalities. Early approaches focused on the joint profiling of gene expression and chromatin accessibility [3] as well as gene expression and surface protein abundance [16]. More recently, methods obtaining both RNA and DNA sequencing from the same single cell have also emerged [17]. Beyond the joint profiling of two modalities, several platforms now support the integration of three molecular layers; for example, Chromatin Accessibility, Interaction and RNA profiles (ChAIR) [18] simultaneously measure gene expression, chromatin accessibility and chromatin conformation. Furthermore, Combined Assay of Target Chromatin Indexing and Tagmentation (COTACIT) [19] simultaneously quantifies H3K27ac, H3K27me3 and H3K9me3 histone modifications in the same single cell. In many studies [19, 3], gene expression profiles reliably distinguish cell types, yet lineage relationships are often captured only by less frequently profiled modalities. This observation underscores the need for integrative approaches that provide a more comprehensive view of cell state.

As experimental approaches continue to evolve, computational methods have been developed to help characterize these increasingly complex datasets. To integrate molecular modalities and encapsulate the full state of a cell, many existing methods capture multimodal information using variational autoencoders (VAEs) [20]. VAEs embed high-dimensional data into a low-dimensional latent space and subsequently reconstruct input data with a decoder network using the latent space as input. Building upon this general framework, each algorithm introduces its own unique modifications; for instance, to facilitate the integration of unpaired data, multiVI [21] trains a discriminator network to ensure that modality-specific encoders construct a similar latent space. When only one modality is available, the encoder maps the observed data into the latent space, and the decoder of the other modality is then used to reconstruct the unobserved modality. This adversarial training scheme is also adopted in GLUE [22], a multimodal integration method that exploits known connections between modalities (for instance, assuming that the expression of a gene co-varies with the openness of the chromosome on which it resides) to facilitate data integration. However, adversarial discriminator networks are known to be unstable and computationally expensive to train [23, 24]. Improving upon this approach, multigrate [25] aligns modality-specific latent spaces by minimizing the distance between their probability distributions.

An implicit assumption made in these methods is that the latent spaces they construct capture the true underlying cell state. However, this assumption may break down when measured modalities capture distinct aspects of cell state, a phenomenon we term structural mismatch. Structural mismatch refers to the situation in which the neighborhood structure of a cell differs depending on which molecular modality is used to define it: in situations with structural mismatch, a cell’s nearest neighbors in the space of one modality are not the same as its nearest neighbors in the space of another modality. The existence of structural mismatch between two modalities is direct evidence that each modality captures aspects of cellular state that the other cannot. In other words, structural mismatch is precisely what makes multi-modal measurements scientifically valuable in the first place. In such cases, forcing all modalities to conform to a single latent space may obscure modality-specific signals and misrepresent biological variation. To overcome this issue, we developed Move BeTween modAlities (MBTA), a flexible deep learning framework that allows seamless translation between any number of molecular modalities in single cells. Rather than forcing all modalities into a shared representation, MBTA constructs a separate latent space for each modality and connects them via flow matching without requiring the two modalities to share a common geometry. Through extensive benchmarking, we demonstrate that MBTA reaches state of the art performance under various metrics. When investigating reasons explaining MBTA’s superior performance, we found that the extent of structural mismatch is anti-correlated with the performance of existing algorithms, thus enabling co-embedding methods to perform reasonably well for modalities spanning similar high-dimensional spaces, but worse for modalities with structural mismatch. Finally, leveraging MBTA’s ability to seamlessly integrate multiple modalities, we demonstrated its ability in identifying hidden lineage structure in human breast tumor samples and evolving eight distinct epigenetic modifications during mouse embryonic development. In sum, MBTA enables imputation of unseen single-cell modalities and temporal evolution of multi-modal datasets across time.

## Results

### Structural mismatch in multi-modal datasets

To develop a framework for translating between single-cell data modalities, we first investigated the topological properties of co-profiled datasets. As a conceptual example, consider a simplified system consisting of a single chromatin region and a single gene. In this setting, heterochromatin formation suppresses transcription, while the absence of transcription does not necessarily indicate the presence of heterochromatin (Figure 1A). Thus, active transcription provides strong evidence for an open chromatin state, while transcriptional silence alone is insufficient to determine chromatin accessibility.

**Figure 1:**
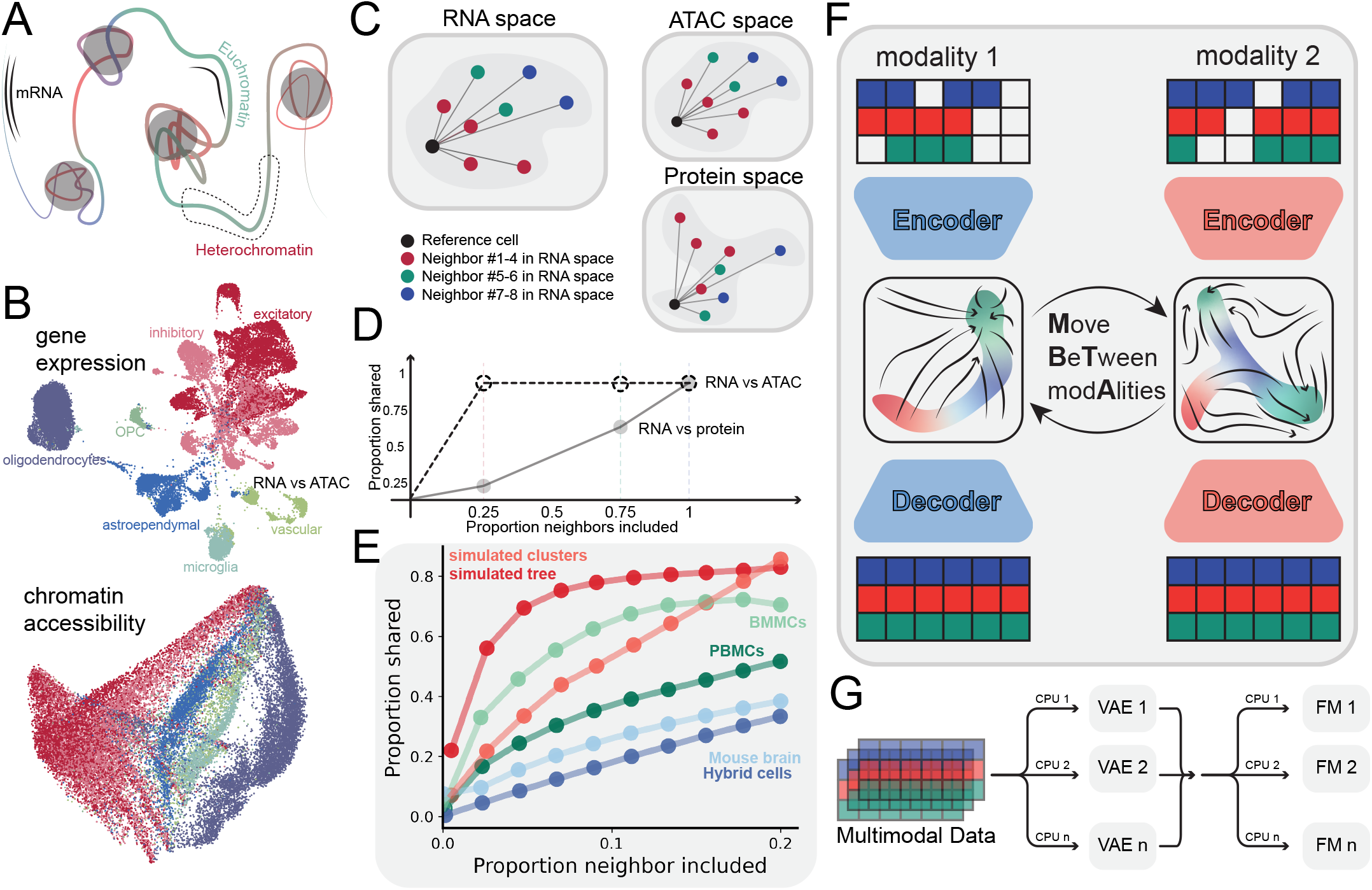
Move BeTween modAlities (MBTA) motivation, design and applications. A: Illustration of a chromosomal segment containing open and closed chromatin. Expressed regions imply open chromatin, but in turn, open chromatin does not necessarily imply expression at that locus. B: UMAP projection of a dataset of mouse brain cells with co-profiled gene expression and chromatin accessibility. C: A reference cell (black dot) and its seven nearest neighbors in RNA space, grouped by rank: neighbors 1–4 (red), 5–6 (green), and 7–8 (blue). In the ATACseq space, the rank ordering of neighbors closely mirrors that of the RNAseq space, indicating similar neighborhood structure. In the protein profiling space, the neighbors identified are different from those in the RNAseq space, indicating structural mismatch between the RNA and protein modalities. D: The proportion of neighbors shared between the RNA space and the ATAC space (dashed) or protein space (solid), plotted as a function of the proportion of neighbors included. The RNAseq–ATACseq neighbor overlap remains consistently high across all neighborhood sizes, whereas the RNAseq–protein overlap increases only incrementally and never reaches a comparable level, quantifying the degree of structural mismatch between these two modalities. E: Relationship between the proportion of neighbors used for the construction of KNN graphs (x-axis) for each modality and the magnitude of overlap of this metric between modalities (y-axis) for the datasets [26, 18, 21, 25] discussed in the main text. F: Overview of MBTA. Separate variational autoencoders (VAEs) are used to encode each modality, and a pairing model trained by flow matching is used to connect the two modality-specific latent spaces. G: Illustration of the MBTA training scheme, where each VAE is trained independently of each other in parallel, with a similar approach for the flow matching models (FMs).

We then investigated whether this asymmetry is reflected in real single-cell datasets. Using paired RNA-seq and ATAC-seq profiles from mouse brain cells [18], we constructed low-dimensional representations of each modality using UMAP [11]. The gene-expression manifold formed distinct, well-separated clusters, whereas the chromatin-accessibility manifold exhibited a more continuous structure (Figure 1B). These differences indicate a structural mismatch between the modalities: cells that are neighbors in the RNA-seq manifold are not necessarily neighbors in the ATAC-seq manifold. Extending this analysis across multiple datasets revealed that the degree of structural mismatch varies substantially across both cell types and molecular modalities (Figure S2A-E). Together, these findings demonstrate that different molecular modalities capture complementary, rather than equivalent, structural representations of cellular state.

To assess the extent of such asymmetries between modalities, we developed a metric to quantify similarity between different modalities co-profiled across single cells. For each modality, we constructed *k*-nearest neighbor (KNN) graphs over a range of *k* values and quantified the proportion of shared neighbors with its co-profiled counterpart (Methods). A simplified example is shown in Figure 1C, where the nearest neighbors of the reference cell in the RNAseq and ATACseq spaces are identical; however, this is not the case for the RNA and protein spaces. As a result, the proportion of neighbors of the reference cell shared between the RNA and ATAC spaces remains high across a range of neighborhood sizes, while the proportion shared between RNA and protein increases more gradually (Figure 1D). Across a diverse collection of co-profiled datasets [18, 21, 25, 26], we observed substantial variability in cross-modality similarity (Figure 1E; Figure S2A–E). For example, the two simulated datasets with co-profiled gene expression and chromatin accessibility [26] (Methods) showed strong concordance, with nearly 80% of the closest 20% of neighbors shared between modalities. In contrast, only about 20% of neighbors were shared across modalities in the mouse brain dataset with co-profiled gene expression and chromatin accessibility [18]. Finally, in a CITE-seq dataset sampled from human bone marrow mononuclear cells (BMMCs) [16], the similarity between modalities was about 60% (Figure 1B).

### The Move BeTween modAlities (MBTA) algorithm

To accurately account for structural mismatches between molecular modalities and faithfully represent cell state, we developed Move BeTween modAlities (MBTA) (Figure 1F). MBTA transforms one modality into another by learning a nonlinear mapping between their latent representations. To capture biological nonlinearity, we parametrize this mapping with a neural network, where each layer introduces additional complexity. When the modalities differ substantially, accurately modeling the transformation of one data modality into another may require a deep network, leading to high memory demands and long training time. To overcome this limitation, we represent the system as a neural ordinary differential equation (ODE) – in effect a neural network with infinite depth [27, 28]. To avoid the computational cost of integrating ODEs during training, we learn the expected gradient directly from data via flow matching [29]. We further reduce computational cost by projecting each modality into a low-dimensional space using a variational autoencoder (VAE) [20] prior to training the pairing model. Thus MBTA constructs modality-specific latent spaces (*z*_1_ and *z*_2_) and bridges them via a flow field, *f_z1→z2_*(*z*_1_*, z*_2_*, t*), such that solving the differential equation 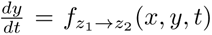 yields the corresponding latent coordinate in *z*_2_ for a cell whose latent coordinate in *z*_1_ is *x* (Figure S3A, Methods). Although the flow matching component generates a continuous trajectory in the latent space, this trajectory only represents the sampling process and is not biologically interpretable as MBTA is not designed to generate biologically meaningful transitional states. Supplementary Note 1 provides additional explanations regarding the flow matching component of MBTA.

MBTA offers several advantages beyond imputing unmeasured modalities. First, it can be trained on datasets where missingness is structured by cell type, where some cell types are fully co-profiled and others are not, differing significantly from scenarios where only a subset of cells is co-profiled. By learning the relationships between modalities from co-profiled cell types, MBTA can generalize this knowledge to bridge modalities in cell types lacking paired measurements (Figure S3B). Second, MBTA can construct a cell-like object, where induced changes in one modality can lead to changess in the other modality. Leveraging this ability, MBTA can quantify the extent of inter-modal coupling; for instance, it can estimate the expected change in protein abundance associated with a change in gene expression (Figure S3C). By analyzing how changes in one modality relate to changes in another across many cells, MBTA is able to identify cell types where modalities become decoupled. Overall, MBTA’s design allows it to operate effectively in realistic datasets with cell type-dependent co-profiling and to generate hypotheses by examining cells with highly or weakly coupled molecular modalities, as we later demonstrate (Figure 4).

In contrast to existing approaches [25, 21], MBTA constructs a separate latent space for each modality and subsequently connects them via flow matching. This architecture insulates each latent space from structural mismatch, an issue that becomes more pronounced as more modalities are integrated. By maintaining modality-specific latent spaces, MBTA also enables each latent space to scale appropriately with its input dimensionality. For instance, projecting modalities with vastly different dimensions (e.g., 50 vs. 1 million) into a shared latent space can under-represent the high-dimensional modality. MBTA’s design can accommodate such disparity with a tailored embedding dimension. Crucially, the modular design of MBTA allows every component to be trained independently: VAEs are fitted to their respective modalities, and subsequently, flow matching models are trained on modality pairs without modifying the underlying VAEs. The entire framework is therefore fully parallelizable, making it possible to integrate an arbitrary number of modalities at the computational cost of integrating just two (Figure 1G).

### MBTA reconstructs and translates accurately

To evaluate the performance of MBTA in reconstructing and translating data, we performed benchmark analyses comparing its performance to that of other multimodal approaches such as multigrate [25], multiVI [21] and GLUE [22] as well as unimodal approaches such as scVI [30]. To this end, we used diverse co-profiled datasets encompassing measurements of gene expression, chromatin accessibility, surface protein abundance and copy number alterations. Specifically, we benchmarked all methods on a hybrid cell dataset [18], peripheral blood mononuclear cell (PBMC) [21] dataset and simulated [26] data with co-profiled gene expression and chromatin accessibility; a hematopoetic stem and progenitor cell (HSPC) [31] dataset measured by CITE-seq; and ER+ breast cancer samples [17] co-profiled using RNAseq and DNAseq. Supplementary Note 2 provides runtime analysis and details on the hyperparameter choices while Figure S1 provides details on the normalization scheme. Each dataset was analyzed to obtain a “data triplet” consisting of the observed, model-reconstructed and model-translated data of the same modality (Figure 2A). The data triplets were co-embedded via UMAP to qualitatively and quantitatively assess the performance of each method (Figure 2B, C). Across multiple datasets, we observed that tested algorithms, with the exception of MBTA, cannot generate data triplets that show large overlap when co-embedded; based on UMAP visualizations, multigrate performed well in data reconstruction but less consistently in translation, whereas multiVI typically generates data with systematic differences with respect to the observed data (Figure 2D-G, Figure S4 and Figure S5).

**Figure 2:**
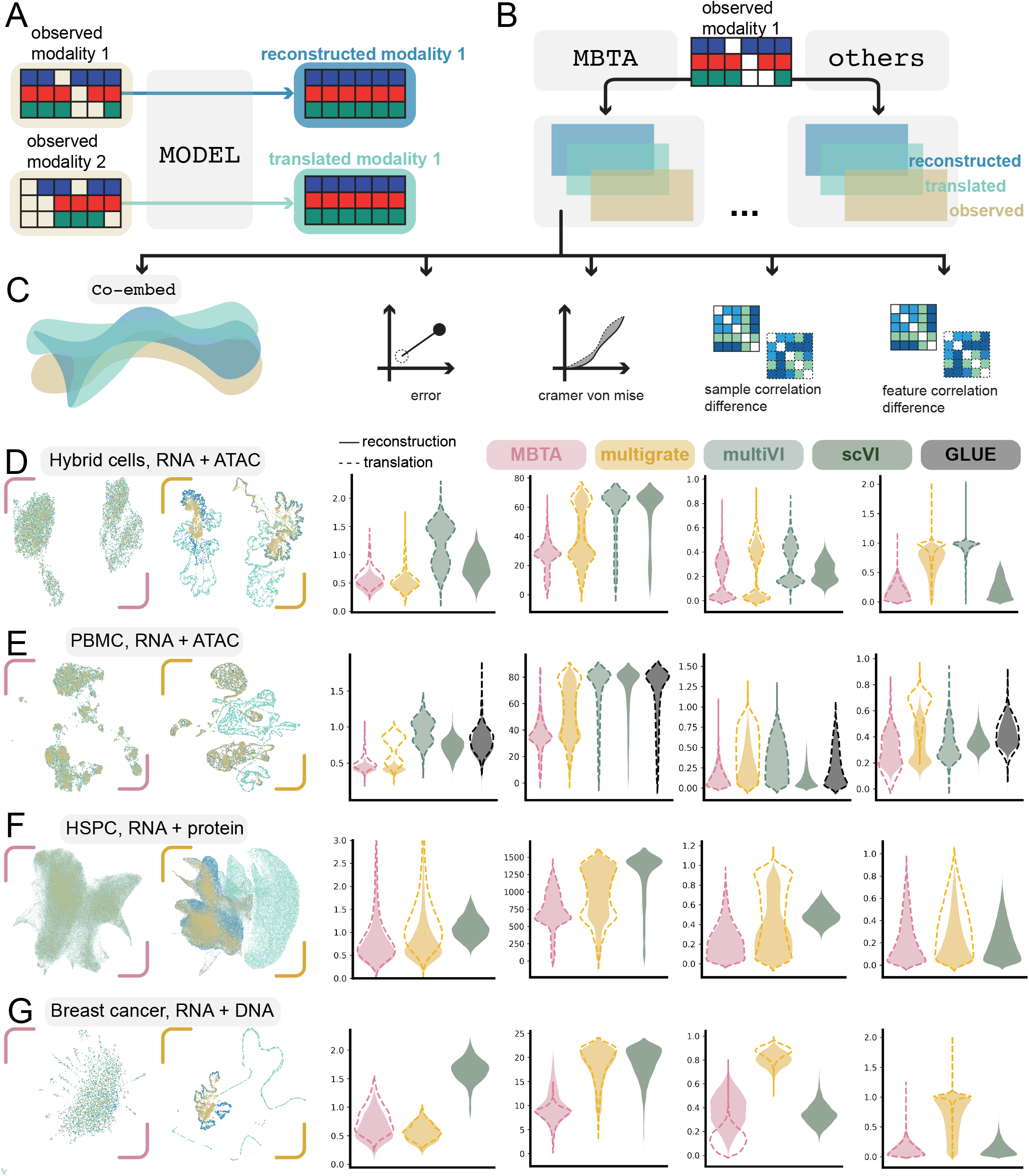
Evaluation of MBTA’s performance. A: A data triplet consisting of the observed modality 1, reconstructed modality 1 obtained from the model by inputting the observed modality 1, and translated modality 1 obtained from the model by translating modality 2 into modality 1. A similar data triplet can be obtained for modality 2. B: Data triplets are constructed using MBTA and comparator methods. C: Qualitative and quantitative measures used to evaluate the tested methods. D-G, results obtained from the hybrid cells [18] (D), PBMC [21] (E), HSPC [31] (F), and ER+ breast cancer [17] (G) datasets.

These qualitative results were then verified using a set of quantitative, orthogonal metrics including mean squared error (MSE), Cramér–von Mises (CvM) statistics, and correlation preservation (Figure 2C). We found that multiVI had the largest mean squared error (MSE) among the methods tested and MBTA performed similarly to multigrate in terms of data reconstruction (Figure 2D-G and Figure S5) but significantly outcompeted multigrate in almost all cases for data translation (Figure 2D-G and Figure S5). To measure how well reconstructed and translated data agree with the observed data on the population level, we computed the CvM statistics to quantify the distance between the cumulative distribution function of the observed and the reconstructed/translated data on a per-feature basis. In all tested datasets, we observed that MBTA had the lowest CvM statistics both in reconstruction and translation (Figure 2D-G and Figure S5); in contrast, multiVI generated data with the largest CvM statistics. This result suggests that in scenarios in which MBTA and other approaches have a comparable MSE, the modalities MBTA reconstructs or translates provide a better match to the observed data on the population level.

To assess each method’s ability to preserve relationships between samples and features, we measured the difference between sample–sample and feature–feature correlations in the original versus reconstructed or translated data. The sample correlation difference reflects whether cells with similar expression profiles remain similar after reconstruction, and the feature correlation difference assesses whether correlated features retain their correlation after model processing. A model that reconstructs or translates perfectly would yield a correlation difference of zero. For data reconstruction, MBTA achieved the lowest sample-level correlation difference among all multiomic methods (Figure 2D–G, Figure S5). In contrast, multigrate’s performance deteriorated substantially during data translation. Feature-level analyses supported this finding, with the feature-feature correlation differences for both multigrate and multiVI deviating from zero (Figure 2D–G).

To understand why existing methods display diminished performance in this setting, we turned to scVI — the unimodal counterpart of multiVI — as a diagnostic reference. Because scVI reconstructs a single modality without any cross-modal alignment, it provides a natural baseline for the correlation structure achievable in the absence of structural mismatch, and allows us to isolate whether multiVI’s underperformance stems from its reconstruction capacity or from the constraints imposed by its shared latent space. We found that scVI performed similarly to MBTA and markedly better than multiVI and multigrate (Figure 2D–G, Figure S5). Since scVI shares the design of the neural network architecture and training scheme of multiVI but operates on only a single modality, its robust performance suggests that the performance reduction of multiVI does not stem from limitations in its reconstruction capacity, but rather from the constraints imposed by forcing modalities with incompatible neighborhood structures into a shared latent space. This finding may generalize beyond multiVI and multigrate, and points to a deeper structural problem with unified latent space approaches as a class. Any method that constructs a shared representation across modalities must resolve a fundamental tension between modality-specific geometries and a single shared coordinate system. Importantly, this shared latent space assumption introduces a compounding problem: as the number of modalities increases, the probability that all modalities share a compatible latent geometry decreases. Each additional modality brings its own neighborhood structure, and forcing all of them into some aligned latent space means that structural mismatch — already demonstrated to be a real and consequential phenomenon in the two-modality case — accumulates with every modality added. The result is a latent space that is increasingly compromised by competing geometric constraints, none of which are explicitly modeled or corrected for. MBTA sidesteps this accumulation entirely and, as shown above, does so without any loss in the correlation preservation that scVI achieves in the single modality case.

### Structural mismatch sets a fundamental limit on model performance

To further evaluate MBTA and the impact of structural mismatch, we analyzed topological metrics that quantify similarity and mismatch across co-profiled modalities. A successful integration method should satisfy two criteria simultaneously (Figure 3A). First, each reconstructed modality should closely match its corresponding observed modality, meaning that intramodal similarity should be high. For example, the nearest neighbors of a cell in the reconstructed gene expression space should correspond closely to its nearest neighbors in the observed gene expression space, and similarly for chromatin accessibility. Second, the similarity between the two reconstructed modalities should reflect the similarity between the two observed modalities. Because structural mismatch is an intrinsic property of the data, a faithful reconstruction should preserve it: reconstructed modalities should not become more similar to each other than the observed modalities are. The observed intermodal similarity (black dashed line, Figure 3B) therefore provides a natural baseline that a well-performing method should reproduce. An ideal method thus yields intramodal similarity curves (colored lines, Figure 3B) that are as high as possible, while its intermodal similarity curve follows the observed baseline (black dashed line, Figure 3B).

**Figure 3:**
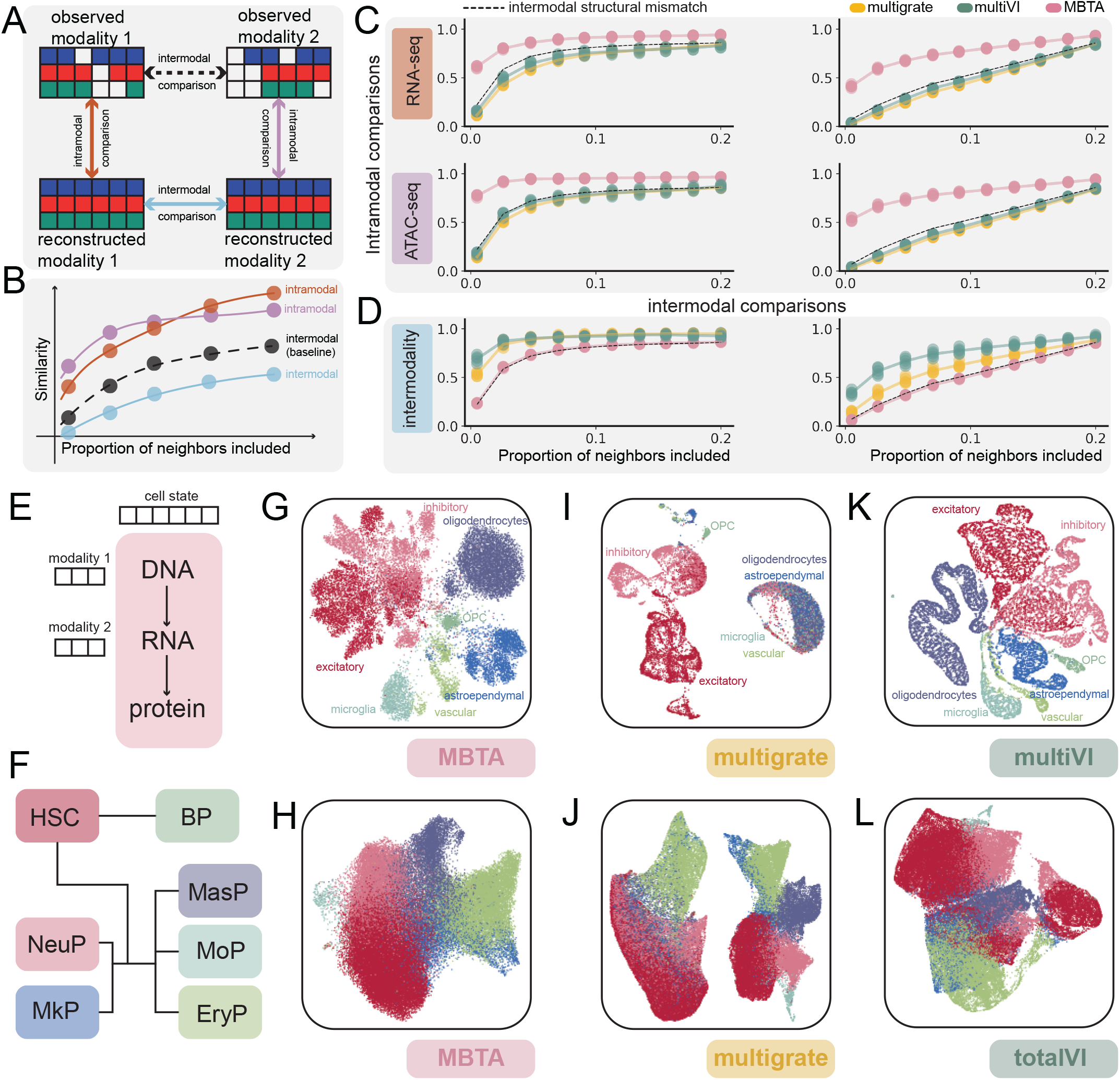
Structural mismatch sets a fundamental limit to model performance. A: Schematic of the comparison of the KNN graphs. Each double-headed arrow indicates a comparison between observed, reconstructed, or translated data. B: An example output for one single method from the analysis is shown, with the color code as indicated in the top panel. C: Similarity between KNN graphs constructed from the observed and reconstructed gene expression (first row) and the observed and reconstructed chromatin accessibility (second row). C: Similarity between KNN graphs constructed from the reconstructed gene expression and the reconstructed chromatin accessibility. The black dashed line represents the baseline intermodal structural mismatch computed from the observed gene expression and the observed chromatin accessibility. Results are computed from synthetic data with a tree-like (left) and a cluster-like (right) topology. Proportions of neighbors included is the ratio between the sample size and the number of nearest neighbors used for the construction of the KNN graph. E: Illustration of how MBTA concatenates modality-specific latent spaces to construct a cell state representation. F: Lineage structure of hematopoetic stem cell differentiation. G-L: UMAP projection of the cell state representation of a mouse brain dataset and HSPC dataset constructed using MBTA (G, H), multigrate (I, J), multiVI (K) and totalVI (L). Panels G, I and K are constructed using the mouse brain dataset; and panels H, J and L are constructed using the HSPC dataset. totalVI is the equivalent of multiVI for CITE-seq data.

Using the similarity between the two observed modalities as a baseline (Figure 3C, D), we then evaluated MBTA. We found that MBTA-reconstructed modalities display higher similarity to their corresponding observed modalities than this baseline (Figure 3C). This observation indicates that MBTA preserves modality-specific neighborhood structures even in the presence of structural mismatch. Consistent with the expected behavior described above, MBTA also reflects the separation between modalities: the reconstructed modalities maintain distinct neighborhood structures, in line with the differences present in the observed data (Figure 3D).

In contrast, multiVI and multigrate produce less faithful reconstructions of individual modalities (Figure 3C). Consistent with reduced reconstruction fidelity, neither method is able to preserve the differences between modalities. In particular, the reconstructed gene expression and chromatin accessibility profiles generated using multigrate and multiVI exhibit highly similar neighborhood structures, with the degree of similarity between these reconstructed modalities exceeding that observed in the original data (Figure 3D).

Taken together, the results suggest a consistent pattern. For multiVI and multigrate, mapping both modalities into a shared representation reduces the structural differences between modalities, and reduced intermodal separation coincides with less faithful reconstruction of each individual modality. This pattern reflects a tradeoff inherent to approaches that enforce a shared representation: encouraging alignment between modalities can come at the expense of preserving modality-specific structure. In contrast, MBTA avoids this tradeoff by maintaining separate modality-specific latent spaces, and is the only method that simultaneously reconstructs each modality faithfully while preserving the structural differences between them. Similar patterns are also observed across other datasets (Figure S6).

### MBTA accurately captures cell state

To investigate if MBTA’s ability to preserve the structural mismatch between modalities is useful for downstream analysis, we set out to investigate whether MBTA can accurately capture cellular identity across modalities. Since MBTA constructs modality-specific latent spaces, following recent algorithmic advances [32], we assembled a cell state vector by concatenating modality-specific latent space coordinates (Figure 3E). Using a hematopoietic stem and progenitor cell (HSPC) dataset [31] (Figure 3F) and a mouse brain dataset [18] (Figure 3G), containing CITE-seq data and co-profiled gene expression and chromatin accessibility data, respectively, we first investigated MBTA’s ability to classify cellular relationships, observing clear separation of each cell type (Figure 3G, H).

However, when multiVI (or totalVI for CITE-seq data) and multigrate were applied to these datasets, we observed an inability of the multigrate latent space to separate four distinct cell types separable by MBTA (Figure 3I) and an inability of multiVI to distinguish subpopulations of vascular cells (Figure 3K). The consequence of structural mismatch was the most pronounced when totalVI and multigrate were applied to the HSPC dataset; this dataset contains a gene expression space with a tree-like like structure (Figure S7A) that is meaningfully different from its protein abundance space (Figure S7B). When this dataset was used to train multigrate or totalVI, we found that these methods attempt to reconcile these two incompatible neighborhood structures by projecting both modalities into a single shared latent space. Because no single geometry can simultaneously honor both neighborhood structures, the model is forced to compromise: the resulting latent space is distorted, and each annotated cell type becomes artificially subdivided into multiple clusters (Figure 3J, L). These artificial subdivisions are present in neither modality individually and represent model-induced artifacts rather than biological signal. These observations were also quantitatively verified, demonstrating that multigrate in general showed poor performance in separating annotated cell types (Figure S8A, B). To investigate the presence of the two distinct clusters of cells in the multigrate latent space, we compared the KNN graphs constructed using the latent, RNA and protein spaces and observed that one such cluster defines a neighborhood structure similarly to its corresponding protein space, and the remaining cluster defines a neighborhood structure similarly to its corresponding RNA space (Figure S7C, D and E).

Together, our results confirm that structural mismatch between modalities impairs model performance and is consistent with our observation that unimodal approaches often outperform multimodal counterparts with identical neural network architectures. This failure mode reveals a deeper issue with the shared latent space paradigm. A single joint embedding implicitly assumes that there exists a unified low-dimensional representation that faithfully captures the structure of all modalities simultaneously. When structural mismatch is present, this assumption is violated by definition. The model is not simply failing to find the right embedding — it is being asked to solve an ill-posed problem. Forcing a shared representation onto structurally mismatched modalities does not resolve the mismatch; rather, it erases it, destroying the modality-specific information that motivated the multimodal measurement in the first place. As a consequence, projecting mismatched modalities into a common space can distort reconstructions, disrupt correlation structure, and misrepresent cell states. By contrast, MBTA overcomes these limitations, accurately reconstructing and translating modalities while preserving the underlying data topology (Figure 3G, H).

### MBTA predicts unseen cell pairings

To assess whether MBTA can capture biological structure in a data-driven manner, we evaluated its ability to extrapolate to out-of-sample cell pairings (Figure S3B). We considered a setting in which only a subset of cells is co-profiled (Figure S3B) and asked whether MBTA can still link cells across modalities when explicit pairing information is incomplete. In this setting, the VAE component has access to the full dataset, whereas the flow matching component is trained only on the co-profiled subset. We used a bone marrow mononuclear cell (BMCC) dataset [25] with co-profiled gene expression and surface protein abundance, training modality-specific encoders and decoders on all cells. In contrast, the flow matching component was restricted to a subset of cells, thereby simulating the absence of complete cross-modal pairing.

To further increase the difficulty of this setting, each subset was constructed to exclude one annotated cell type, resulting in multiple MBTA models in which pairing information for a specific cell type is entirely missing from the flow matching component. As a control, we trained a baseline model with access to all pairing information, enabling a direct comparison between the fully paired setting and MBTA under incomplete pairing, and allowing us to evaluate its ability to reconstruct out-of-sample cross-modal relationships.

To evaluate the performance of MBTA, we encoded gene expression into the RNA latent space and translated it into the protein latent space. Next, for cells with unknown pairing, we quantified their distance to the centroid of each labeled cluster. Successful extrapolation would place projected coordinates close to their originating cluster (in-group) and not with any of the other clusters (out-group). Indeed, we found that the in-group distances were always lower than their outgroup counterparts, although the extent of their difference varied by cell type (Figure 4A), with similar results obtained when protein abundance was used as input (Figure S9A). Using these distances for cluster assignment, MBTA achieved an average area under the receiver operator characteristic curve (AUC) of 0.88 across cell types, where 15 out of the 24 cell types had an AUC over 0.9 (Figure S9B, and Figure S10A). Comparison with the baseline model confirmed that full pairing information yields higher accuracy (Figure S10B and C). Furthermore, we visually confirmed the ability of MBTA to conduct out-of-sample pairing via co-embedding the observed and translated protein abundances of unseen cell types (Figure 4B). Collectively, our findings demonstrate MBTA’s capacity to extrapolate out-of-sample pairings by leveraging information from co-profiled samples.

**Figure 4:**
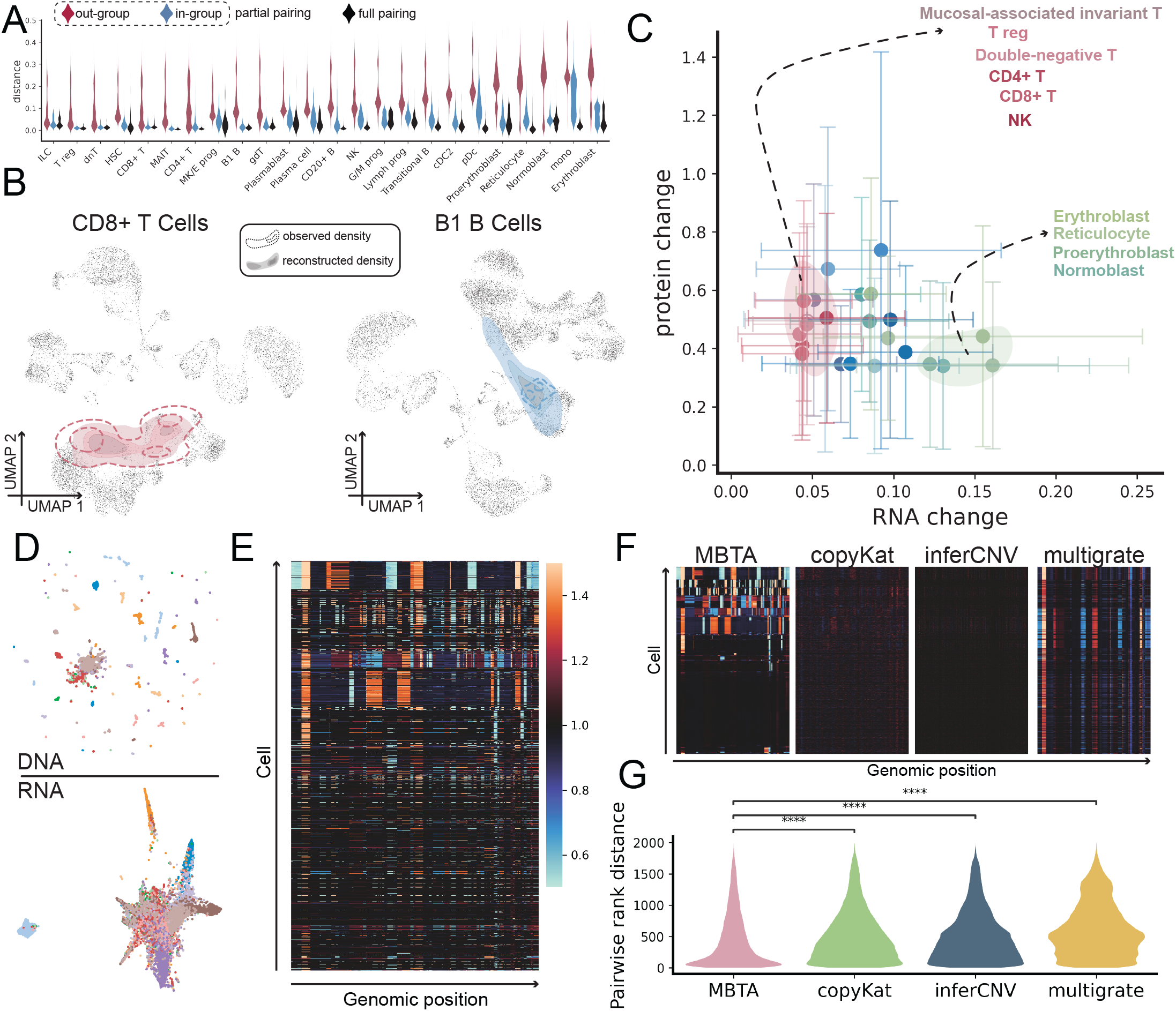
Applications of MBTA. A: MBTA leverages known pairing information from a subset of BMCC cell types to infer the pairing of unseen cell types. Distribution of the distance between each cell to the centroid of the cell type it belongs to (in-group) and all remaining cell types (out-group) when the pairing information between a said cell type (x-axis) is unknown to the model (partial-pairing) as well as results computed from a model with all pairing information (full paring). These distances are used to predict the originating cell type of each cell and the results are quantified via the receiver operating characteristic curve, results shown in Figure S10A. B: UMAP projection of the predicted distribution of CD8+ T cells (left) and B1 B cells (right), remaining cells shown in black as background. C: Relationship between RNA change and protein change as quantified by MBTA’s perturbation-based coupling analysis in a set of BMCCs. Each circle represents one cell type, with the horizontal position indicating the mean change in gene expression and the vertical position indicating the mean change in protein abundance in response to a random Gaussian perturbation applied to the RNA latent space. Error bars show the standard deviation across repeated perturbations for each cell type. Colors denote cell type identity. The broken arrows point to cells belonging to the T-cell (red) and erythrocyte lineage (green). D: UMAP projection of the observed gene expression and copy number profile. E: The ground truth copy number profile of test cells randomly selected from 12 ER+ breast cancer patients, with colors indicating copy number state. F: The four heatmaps show the copy number profiles reconstructed by MBTA, copyKat, inferCNV, and multigrate respectively, plotted on the same color scale as the ground truth to allow direct visual comparison. The color bar is shared with panel E. G: The violin plots quantify the pairwise rank distance error. For each method, pairwise distances were computed in the predicted and observed copy number spaces, converted to ranks, and the absolute difference in ranks is displayed as the error for each cell pair. A lower distance indicates that the predicted profiles preserve the relative copy number relationships between cells more faithfully. For panel G, *p* values are derived from a two-tailed Mann-Whitney U-test. \*\*\*\**p <* 0.0001.

### MBTA quantifies the strength of modality coupling

Encouraged by MBTA’s ability to bridge across modalities, we next developed an analysis pipeline to quantify the coupling strength between modalities by measuring how perturbations in one modality propagate to the other (Figure S3C). Briefly, we first applied a small perturbation to the latent space of modality 1. We then used MBTA to map both perturbed and unperturbed latent representations of modality 1 data into the space of modality 2, and decoded them back to the original feature space. This approach allows quantification of the changes in modalities 1 and 2 in response to a controlled, finite perturbation in modality 1.

Applying this analysis to the BMCC dataset [25], we found that changes in gene expression are not systematically associated with changes in protein abundance (Figure 4C), despite this dataset exhibiting relatively strong overall concordance (Figure 1E). Instead, the relationship between modalities varies across cell states. For example, in T cells, protein surface markers show larger changes than gene expression, while in erythropoietic populations (erythroblasts, reticulocytes, and erythrocytes), the opposite pattern is observed. The behavior in T cells is consistent with the relatively modest correspondence between surface protein abundance (e.g., CD4, CD8) and mRNA levels [16], and with known regulation of these proteins through post-translational mechanisms [33, 34, 35, 36, 37]. In contrast, the stability of protein levels in erythroid-lineage cells is consistent with the well-characterized process of organelle clearance during erythropoiesis, including loss of ribosomes [38, 39].

Consistent with these patterns, direct visualization of the data manifold shows that gene expression separates distinct erythroid populations, while their surface protein profiles are largely indistinguishable (Figure S10D). Together, these results suggest that MBTA can identify regions of modality-specific decoupling in gene expression or protein space, which manifest as local structural mismatches between modalities.

### MBTA reveals hidden tumor lineages

We next sought to investigate whether MBTA is capable of inferring tumor clonal structure from scRNA-seq data. To this end, we leveraged recent technological advancements allowing gene expression and copy number profiles to be simultaneously measured from the same single cells sampled from 12 ER+ breast tumor patients [17]. Lineage relationships in this dataset can be ascertained either using the DNA sequencing data, enabling direct determination of copy number alterations (CNAs), or using the RNA sequencing data, from which CNAs can be inferred by employing computational pipelines such as inferCNV [40] or copyKat [41]. The structural mismatch between the RNA and DNA modalities is apparent when inspecting UMAP visualizations of each modality (Figure 4D and Figure S11). To investigate whether MBTA is able to accurately translate between these modalities with pronounced structural mismatch, we used a subset of the co-profiled cells to train MBTA and used the remaining cells to evaluate its performance. We also implemented the same training-test pipeline using multigrate. Finally, we applied inferCNV [40] and copyKat [41] to the test cells to infer CNAs from scRNA-seq data and compared the inferred CNA profiles to the ground truth of DNAseq-called CNAs (Figure 4E). We observed that MBTA was the only method able to detect clear chromosomal gains and losses (Figure 4F) and has the lowest error when evaluated using ground-truth data (Figure 4G). This performance difference reflects a fundamental difference in modeling assumptions: both inferCNV and copyKat assume that the expression level of a gene is entirely explained by its own copy number, ignoring gene–gene interactions. MBTA, by contrast, makes no such assumption and learns the relationship between gene expression and copy number directly from the data. In situations where gene expression is not entirely determined by DNA copy number at the same locus, MBTA’s lack of restrictive assumptions allows it to outperform methods that rely on this simplification. Indeed, numerous studies have ascertained that gene expression in aneuploid tumors can be buffered [42] and even uncoupled [43], demonstrating that gene expression and copy number can vary independently.

The capacity of MBTA to integrate distinct modalities and enable precise cross-modal transitions allows investigation of inter-modality relationships that were previously inaccessible. Within this framework, our previously established coupling analysis (Figure S3C) can be conceptualized as a series of in silico perturbation experiments. In each experiment, a minor alteration is introduced into a cell’s transcriptomic state, and MBTA is used to infer the corresponding modification in its copy number profile required to achieve the specified change in gene expression. This framework enables systematic characterization of the relationship between gene expression and copy number alterations (CNAs), thereby identifying genes most strongly associated with CNA-driven expression changes and genomic regions most likely to exert transcriptional effects. Applying this framework to co-profiled ER+ breast tumor samples revealed that the genes most sensitive to CNAs include PRLR [44, 45], IGF1R [46, 47], UVRAG [48], and GREB1L [49], with MDK [50, 51, 52, 53] showing the strongest sensitivity. Several of these genes have established roles in ER+ breast cancer; notably, MDK is upregulated in patients [51] and activates the PI3K–AKT–SREBF pathway, a known driver of tumorigenesis [50]. Visualization of cells expressing dosage-sensitive genes showed that they localize to transcriptomic regions characterized by extensive clonal heterogeneity (Figure S12A and B). Given their diverse clonal backgrounds (Figure S12C), our results suggest a possible convergence event during tumor evolution. Moreover, chromosomes 3, 8, 17, and 20 were most strongly associated with dosage-responsive genes (Figure S12D), consistent with the empirical correlation between gene expression and copy number (correlation: 0.52, *p* = 10*^−^*^30^) and with prior experimental observations [54, 55, 56, 57, 58]; for instance, chromosomes 8 and 17 are known rearrangement hotspots frequently linked by interchromosomal translocations [56] and their amplification predicts early relapse in ER+ breast cancer [54]. Together, our analyses reveal that MBTA can be used to infer hidden tumor clonal structures and identify dosage-sensitive genes.

### Temporal modeling of embryonic development with MBTA

Finally, we evaluated MBTA’s performance on a dataset containing more than two modalities, a recently published dataset [19] that jointly profiled H3K9me3, H3K27ac and H3K27me3 in single cells from different stages of mouse embryonic development (Figure 5A, B). We first evaluated MBTA’s performance in constructing a cell embedding for this co-profiled, tri-modal dataset, observing that cells of distinct embryonic stages are well separated in the low-dimensional space (Figure 5C). When benchmarking MBTA’s performance on this dataset against multigrate’s ability to construct a cell embedding (Figure 5D), we found that unlike MBTA, the latter failed to capture true cellular states, producing nearly identical representations for cells from the 2-cell and 4-cell stages. Quantitative analysis based on the proportion of each cell’s top five nearest neighbors sharing its embryonic stage confirmed this observation: while MBTA embeddings correctly identified similar cells across all types, multigrate produced overlapping representations for cells from the 2- and 4-cell stages and further subdivided zygotic cells into artificial subpopulations not present in the original data (Figure 5E). In addition to the co-profiled data, the mouse embryonic development dataset [19] contains unimodal measurements spanning eight modalities across embryonic development (Figure 5B). These measurements include seven epigenetic marks—H3K4me3, H3K4me1, H3K36me3, H3K27ac, H3K27me3, H3K9me3, and H2A.Z—alongside scRNA-seq. Using this octa-modal dataset, we evaluated MBTA’s capacity to model temporal cell evolution (Figure 5F). MBTA was trained to translate each epigenetic modality from scRNA-seq data, with cells randomly paired by embryonic stage during each training iteration to mitigate the absence of co-profiled measurements. To confirm that MBTA’s translation performance is robust to annotation noise, we performed a label-shuffling experiment in which an increasing proportion of cell annotations were randomly permuted prior to training; MBTA retained strong predictive performance even at substantial levels of label corruption in both real and synthetic datasets (Figure S19–S22). Embryonic staging is rarely ambiguous, lending additional confidence to the pairings learned by MBTA. We then applied an optimal transport–based algorithm [59] to construct a flow map linking transcriptomic states. The zygotic transcriptome was evolved forward in time and subsequently translated by MBTA into the seven epigenetic modalities, reconstructing the complete developmental cell state. Across modalities, the predicted future profiles closely matched observations: in all cases, projected zygotic cells were nearest, in geodesic distance, to their corresponding target cells (Figure 5G, S13, S14).

**Figure 5:**
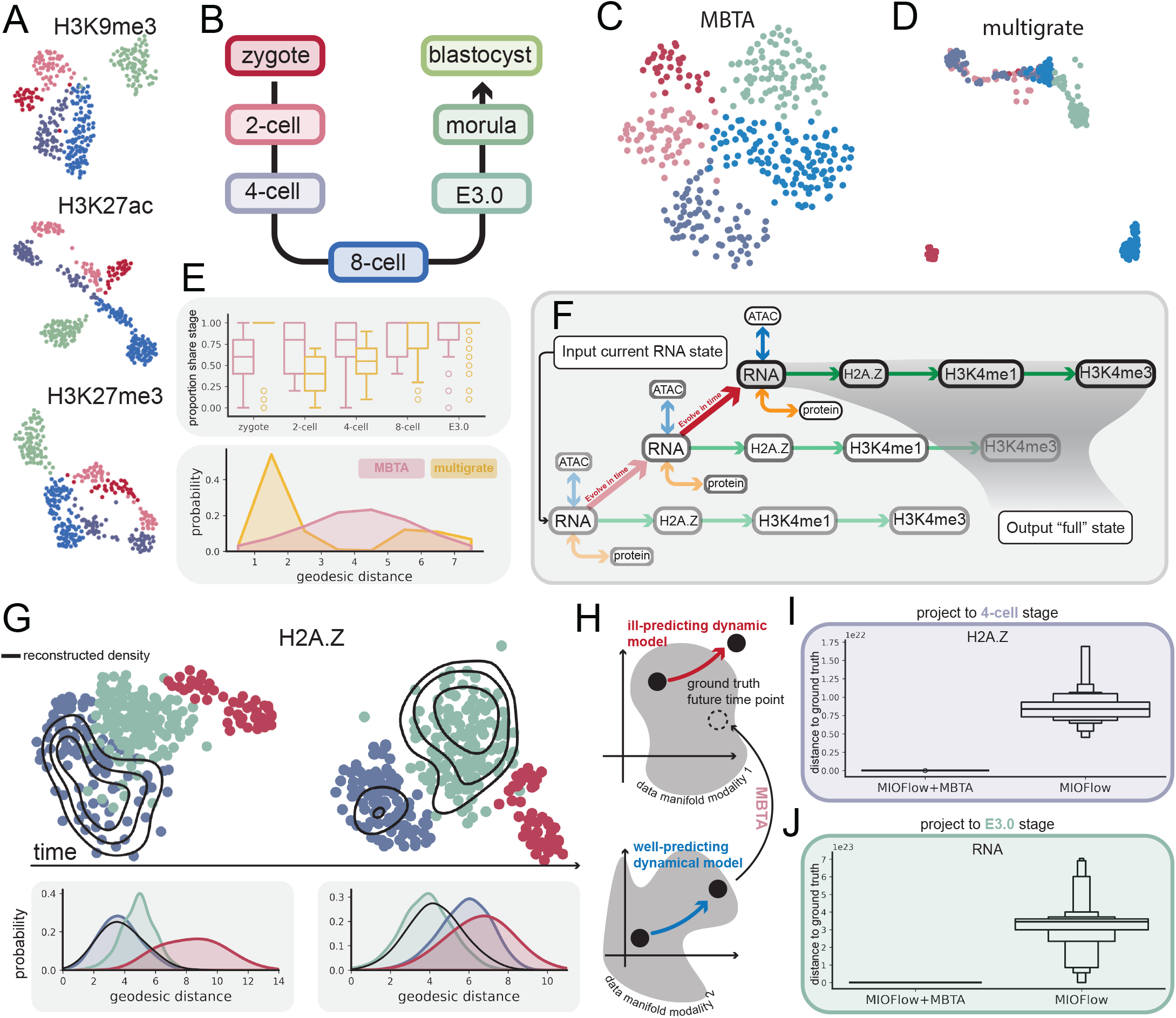
Construction of an evolving embryo. A: UMAP projection of the co-profiled epigenetic features of a developing mouse embryo [19]. B: Schematic of embryonic development profiled in the same study. C: MBTA-constructed cell representation using information from all three modalities shown in A. D: multigrate latent space incorporating information from the same three modalities. E: Proportion of cells around each cell that share its embryonic stage (top) and the distribution of the geodesic distances of all zygotic cells (bottom). F: Illustration of how MBTA contributes to the construction of an evolving embryo. First, gene expression is evolved in time followed by the use of MBTA to translate gene expression to other epigenetic features. G: UMAP co-embedding of the observed H2A.Z and the model constructed H2A.Z (black contour) of cells belonging to the 4-cell (left) and the E3.0 (right) stage. Below each co-embedding, the distribution of geodesic distances between the target cells and cells of each embryonic stage as well as the model-constructed data (black) are shown. H: Cartoon illustrating the principle of the “evolve and translate” strategy when there is an illpredicting model that cannot accurately evolve modality 1. In this situation, MBTA is used to translate the output from the well-predicting dynamical model of the other modality for a more accurate prediction of modality 1. MIOFlow is used to construct a unimodal flow for RNA and H2A.Z data. The boxplot quantifies the distance between model-generated samples and target samples using either the “evolve and translate” strategy or MIOFlow directly to the 4-cell (I) or the E3.0 (J) stages.

MBTA can operate independently of flow maps, enabling any single-modality algorithm to evolve the full cellular state over time. When a modality is difficult to model—due to high manifold curvature or other algorithmic limitations—MBTA allows training on a more tractable modality that can then be translated to others (Figure 5H). To evaluate this “evolve and translate” strategy, we trained unimodal flows for gene expression and H2A.Z dynamics using MIOFlow [60]. We observed that MIOFlow was unable to accurately evolve zygotic H2A.Z and gene expression to the 4-cell stage and E3.0 stage, respectively (Figure 5I, J). For example, we found that predicting the 4-cell H2A.Z data was more accurate when evolving gene expression with MIOFlow and then translating it to H2A.Z via MBTA (Figure 5I). Whenever direct temporal evolution with MIOFlow fails, MBTA can be used to translate the more accurate modality to predict the modality that is more difficult to evolve (Figure 5I, J and Figure S16). Similar results were obtained using TIGON [61] instead of MIOFlow: direct evolution of gene expression produced UMAP embeddings that failed to overlap with the ground truth of the 4-cell stage (Figure S15A), while evolving H2A.Z and translating it to gene expression yielded substantial concordance (Figure S15A). Quantitative analyses confirmed these results (Figure S15B) and similar patterns were observed when zygotic cells were projected to the E3.0 stage (Figure S15C and D).

Together, these results demonstrate that MBTA can seamlessly integrate an arbitrary number of modalities. We further established a proof of principle for coupling MBTA with dynamical systems to create a multimodal virtual cell entity capable of autonomous temporal evolution. Finally, we show that MBTA can enhance existing single-modality algorithms through an “evolve and translate” strategy to improve their predictive performance.

## Discussion

Here we introduce MBTA, a flexible deep learning framework designed for seamless integration of single-cell multi-omics data. In this work, we identified a previously unrecognized phenomenon in multi-omic data: structural mismatch. Structural mismatch refers to differences in neighborhood structure across modalities, where a cell’s nearest neighbors in one modality need not correspond to its nearest neighbors in another. We found that structural mismatch is a prevalent property of multi-omic datasets and, across multiple settings, can substantially limit the performance of existing approaches [21, 25]. Importantly, this phenomenon is not a technical artifact, but a biologically informative signal that reflects the fact that different modalities capture complementary aspects of cellular state. If two modalities were perfectly aligned in their neighborhood structure, they would be largely redundant, and jointly measuring them would provide limited additional information.

MBTA is, to our knowledge, the first method explicitly designed to address structural mismatch between modalities – a problem that has not previously been identified or formally studied in the single-cell multi-omics literature. Instead of enforcing a shared latent space across modalities, MBTA learns separate latent spaces for each modality and connects them via flow matching. When relationships between modalities are not strictly one-to-one, a single state in modality 1 may correspond to multiple plausible states in modality 2 and vice versa. In this setting, flow matching is well suited, as it transports probability mass toward the full conditional distribution rather than collapsing heterogeneous targets into a single averaged representation. As a result, branching structure, fate bias, and rare regulatory configurations can be represented as distinct high-probability regions in the target space, rather than being smoothed into intermediate states. We further found that MBTA’s advantages are not limited to regimes with strong structural mismatch. In datasets where modalities exhibit similar neighborhood structures, such as the synthetic cluster dataset investigated (Figure S2C), performance differences between methods become smaller, as expected. However, even in these low-mismatch settings, MBTA matches or outperforms competing approaches in translation tasks and in preserving both sample-level and feature-level correlation structures (Figure S5A). This observation indicates that the modality-specific latent space design does not incur a penalty in well-aligned data, and instead provides a consistent framework across the full spectrum of structural mismatch – from largely congruent modalities to strongly discordant ones. Together, these results show that MBTA can overcome a fundamental performance limitation imposed by structural mismatch and can be used to extract meaningful cell state representations while jointly modeling the evolution of multiple modalities over time.

Compared to other VAE-based approaches, MBTA holds several key advantages. First, MBTA can be used for non-co-profiled data with cell type labels. To demonstrate this capability, we used MBTA to connect eight modalities measured independently in cells at clearly defined embryonic stages (Figure 5). Second, MBTA is able to learn the pairing principles between different modalities during training and generalize these learnings to cells with unknown pairing (Figure 4A). Third, by constructing modality-specific latent spaces, MBTA allows the dimensionality of each latent space to be chosen independently to match the complexity of its corresponding input modality. A high-dimensional modality such as chromatin accessibility, measured across tens of thousands of peaks, may require a higher-dimensional latent representation to faithfully capture its structure, while a lower-dimensional modality such as a surface protein panel may be well-represented in a much smaller latent space. Together, these abilities of MBTA allow the simultaneous analysis of co-profiled and non-co-profiled datasets and the integration of modalities of different dimensionality, making it a powerful tool to harmonize multimodal and unimodal single cell atlases and uncover latent cellular programs that span transcriptomic and epigenomic landscapes.

Because flow matching assumes that the underlying velocity field is known, the target vector field MBTA seeks to learn remains the same regardless of training initialization and the choice of hyperparameters (Figure A2–A5). However, the performance of MBTA can be significantly impaired when the data is not co-profiled and the available cell type annotation is too coarse. In such cases, the within-label heterogeneity may be too large for the flow matching component to learn a biologically meaningful conditional distribution. However, when annotations are sufficiently detailed — for example, distinguishing cell types, developmental stages, or patient-specific subpopulations — MBTA can learn accurate cross-modal mappings even without co-profiled data. The required level of annotation granularity depends entirely on the biological question MBTA is used to address. Coarse labels can be adequate when the goal is to capture broad differences between cell types: for example, labels such as “T cells” and “B cells” might be sufficient when the investigation concerns lineage-level differences in the immune system, because the within-label heterogeneity among T cells is small relative to the between-label differences separating T cells from B cells. The same labels may become insufficient in other situations, for instance when distinguishing effector from naïve T cells, because the biologically meaningful variation now lies within subcategories of the T cell label itself, and MBTA would require more granular annotations to model it accurately. With sufficiently detailed annotation, MBTA is robust to annotation errors in both real and synthetic datasets (Figure S19–S22).

The design of MBTA allows translation of any modality into any other modality via flow matching. As MBTA’s name is inspired by the Boston subway system (the Massachusetts Bay Transportation Authority), we think about this concept in terms of stations and routes. Each modality is a train station, and each trained flow matching component represents a direct route between two stations. Translating from one modality to another requires traveling along one or more routes (Figure S23A, B and C). The key insight is that every additional step (i.e., every intermediate modality a translation must pass through) can introduce additional error. If the objective is to translate between any pair of modalities with the smallest possible error, the all-to-all design (Figure S23B) is the right choice. Every modality is directly connected to every other modality, so every translation is a single step. This appraoch minimizes error but scales poorly. If the objective is only to generate all modalities, the central design (Figure S23A) is sufficient and far more efficient. One modality — typically the most commonly measured one, such as RNA — can serve as the central hub. Every other modality is connected directly to the central modality which reduces error. Starting from the center modality, this design allows all modalities to be generated without compromising the quality of the generated data. This design achieves such high efficiency by introducing the smallest number of additional steps: any modality can be translated to any other in at most two steps, first to the central hub, then out to the target. When designing the most efficient station layout for a given experimental context, we recommend that direct flow models be prioritized for modality pairs that will be translated most frequently, as direct translation avoids intermediate steps and preserves the highest fidelity. Modalities that are needed less often can be connected to a central hub modality rather than to every other modality directly, reducing the total number of flow models required while maintaining full coverage of the translation network. This hybrid design — direct connections for high-priority pairs, hub-mediated connections for peripheral modalities — maximizes overall efficiency without sacrificing biological resolution for the translations that matter most. Importantly, the modular nature of MBTA means that this layout is never fixed: if a new modality pair becomes relevant as the experimental focus shifts, a direct flow model for that pair can be trained independently and added to the existing network at any time, without retraining the VAEs or any of the previously established flow models. This approach allows the translation network to grow incrementally as new biological questions or modality emerge, rather than requiring a full retraining of the entire model. Beyond increasing computational efficiency via the design of more efficient connection schemes, MBTA’s modular architecture confers a more fundamental scalability advantage: every component can be trained in parallel (Figure 1G). As such, each VAE model and each flow matching model can be trained entirely independently of each other. This approach leads to the wall-clock time required to train an MBTA model with ten modalities being essentially the same as training a model with two modalities — the additional components simply run concurrently rather than sequentially. In practice, MBTA’s scaling behavior is closer to constant than quadratic: the architectural cost grows with the number of modalities, but the computational cost in time does not.

The exact directionality of the designed routes does not necessarily have to be imposed by any underlying biological mechanism – such as chromatin accessibility affecting gene expression – but can simply be determined by the availability of data. For instance, for a given experiment resulting in scRNA-seq data, analysis of ATAC-seq data might also be of interest even though that dataset might not be available. MBTA can be used in such situations to generate ATAC-seq data from RNA-seq data. In addition, MBTA also allows users to use custom embeddings, thereby eliminating the need to train modality-specific encoders. Existing foundation models [62, 63, 64, 65] trained on millions of cells can be suitable replacements for scRNA-seq encoders as they can provide good quality zero-shot embeddings (Figure A6, A7, A8 and A9); however, they are generally not designed to reconstruct input data (Supplementary Table A1) and the speed with which these embeddings are generated can differ greatly, ranging from minutes to days for the same dataset.

By linking modalities together, we are able to learn relationships among those modalities and quantify how changes in one modality affect the other. For example, by linking copy number and gene expression data, we were able to conduct “virtual experiments” to alter the transcriptome of cells, infer the minimal CNA required to realize this change, thereby identifying genes that are subject to copy number changes, and pinpoint potential chromosomal loci that drive gene dosage effects. MBTA’s capacity to integrate arbitrary modalities and propagate changes across them makes it a natural foundation for constructing a virtual cell. Using the evolving embryo data, we demonstrated that the most frequently sampled modality, gene expression, can be evolved over time and, when used in conjunction with MBTA, evolve all remaining modalities. Furthermore, we illustrated that MBTA grants any algorithm built to evolve a single modality in time the ability to evolve all modalities. When more and more modalities become available, MBTA integrates them seamlessly without having to re-train the full model. With enough data, MBTA allows us to input one modality and output all other modalities in order to train a dynamical model with all available data. By learning the rules governing how different modalities evolve in time, we open the door to identifying their regulatory hierarchies, simulate perturbation responses across modalities, and design the best course of action for cell state control.

MBTA can be used in conjunction with scVI [30] to query batch-affected datasets. By projecting the data onto the space of the reference dataset, MBTA can then be used to translate the batch-corrected data. We demonstrated that correction using scVI is sufficient for realistic translation (Figure S17, Figure S18). However, users may instead rely on other batch correction methods [66, 67, 68] to project the query data to the reference data. Potential limitations of MBTA include its inability to be directly applied to .fastq files. Instead, the raw sequencing data first needs to be processed to generate gene count/peak matrices. When analyzing multiple datasets at once, this limitation of MBTA will require input data to be processed following a consistent procedure to mitigate potential batch effects and to ensure that a shared set of genes/peaks is called. In addition, VAEs, based on which modality-specific encoders and decoders are built, are inherently limited in generalization [69, 70]. While MBTA performs well when incorporating additional modalities measured from the cell types the model has been trained on, in situations when measurements on new cell types are performed, re-training of the entire model may become inevitable.

To summarize, we developed MBTA to overcome structural mismatch — a phenomenon prevalent in multiomic datasets that has gone unrecognized yet severely hinders the construction of meaningful cell state representations. Through its novel design, MBTA seamlessly connects an arbitrary number of molecular modalities, learns how they are coupled to one another, and evolves them through time. We believe this positions MBTA not merely as a tool for current multimodal datasets, but as a paradigm for integrating the full spectrum of cellular measurements — from transcriptomics and epigenomics to proteomics and beyond — into a unified, biologically faithful model of the cell.

## Methods

### The MBTA model

#### Constructing modality-specific latent spaces with variational autoencoders

MBTA is composed of two major components: (i) modality-specific variational autoencoders to construct meaningful latent spaces, and (ii) flow matching models to bridge different latent spaces. The encoders and decoders of the VAEs are trained using the ADAM optimizer using a learning rate of 0.0005 such that they minimize the reconstruction error. By default, the encoders and decoders contain four layers of neurons with ReLU activation. We followed an approach employed previously [71] to avoid overcrowding of the latent space.

#### Flow matching across modalities

The second component of MBTA constructs flow fields between any arbitrary choice of two modalities with latent spaces *z*_1_ and *z*_2_ constructed using VAEs trained in the previous step. The expected coordinate of a cell, *y*, on *z*_2_ given its coordinate on *z*_1_, *x*, is the solution of the ordinary differential equation, 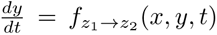, where *f_z1→z2_*(*x*, *y*, *t*) is parametrized by a neural network with four layers of neurons with ReLU activation. For simplicity, we omit *x*, *t* as well as the subscripts in the following sections. The construction of *f* (*y*) begins with the definition of a conditional flow field *f* (*y|y_T_*), which defines the flow to a specific target on *z*_2_, *y_T_* . To obtain the marginal flow field, *f* (*y*), we compute the weighted average of the conditional flow field:

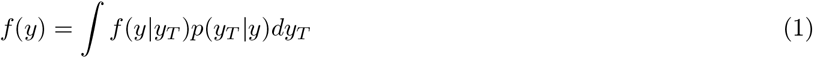

The second term on the right-hand side, *p*(*y_T_ |y*), denotes the probability of a particle to arrive at *y_T_* when subjected to the conditional flow, *f* (*y|y_T_*), given that its current position is *y*. Intuitively, *f* (*y*) should “point to” directions, *f* (*y|y_T_*), that are associated with the largest *p*(*y_T_ |y*), as those are the directions that are the easiest to flow toward given the current position. In practice, however, it is difficult to compute *p*(*y_T_ |y*). By invoking Bayes’ rule, equation 1 can be re-written as:

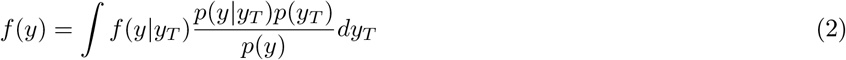

An intuitive way to train a neural network, *f*^^^(*y*), is to minimize

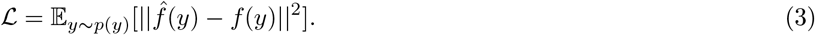

However, the integrand in equation 2 is intractable. To deal with this problem, *L* can be expanded:

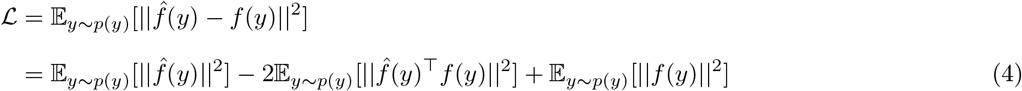

In the above expression, the second term on the right-hand side can be further expanded:

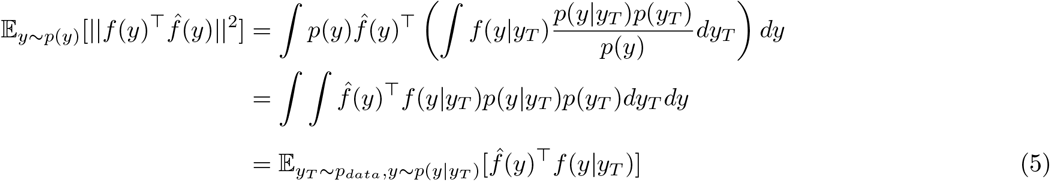

Equation 5 states that E*_y∼p_*_(_*_y_*_)_[*||f* (*y*)*^⊤^f*^^^(*y*)*||*^2^] can be expressed in terms of the expectation of the dot product between the marginal and the condition flow field. However, instead of drawing *y* directly from *p*(*y*), the distribution of *y* subjecting particles to the flow defined by *f* (*y*), we draw *y* by first drawing *y_T_* directly from data, and then subject particles to the flow defined by the conditional flow field *f* (*y|y_T_*). Plugging equation 5 back to equation 4 yields:

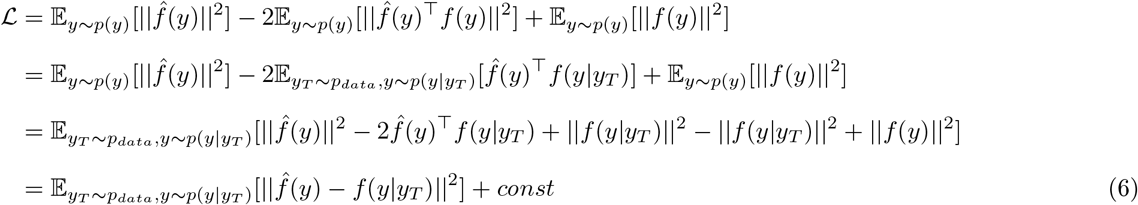

For simplicity, we have chosen to use the Gaussian conditional probability path, defined by:

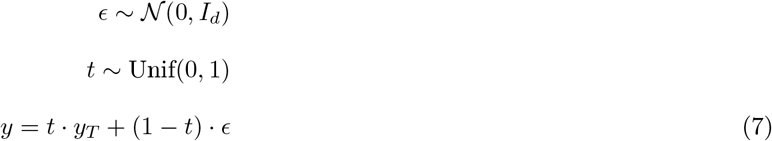

which is associated with the conditional flow field [72, 29]:

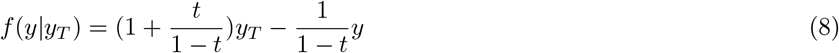

Together, we arrive at our final loss function:

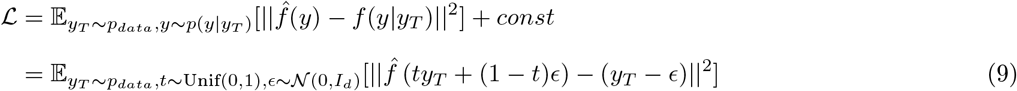

The final loss function aims to construct a flow field, *f*^^^(*y*), that points to data points *y_T_* as expected. Recall that the full form of *f*^^^(*y*) is *f*^^^*_z_*_1_ *_→z_*_2_ (*x, y, t*). During training, co-profiled datasets allow us to directly use the corresponding *x* of each *y*. In situations when we do not have co-profiled data but are provided cell type annotations, we use the provided annotation to connect *x* to *y*.

### Application of existing methods

#### multiVI

multiVI [21] is applied following its tutorial on https://docs.scvi-tools.org/en/stable/tutorials/notebooks/multimodal/MultiVI_tutorial.html. We used the latent space extracted using the model.get_latent_representation() function to represent cell state. To translate from modality 1 to modality 2, we set modality 2 to zero and then used either the model.get_normalized_expression() or model.get_accessibility_estimates() function to generate the missing modality.

#### totalVI

We used totalVI [73] to analyze CITE-seq datasets following https://docs.scvi-tools.org/en/stable/user_guide/models/totalvi.html. Cell state representations are extracted using the model.get_latent_representation() function.

#### scVI/peakVI

To investigate if structural mismatch between modalities impairs model performance, we also used scVI/peakVI to analyze scRNA/scATAC-seq data following tutorials on https://docs.scvi-tools.org/en/stable/tutorials/notebooks/scrna/harmonization.html and https://docs.scvi-tools.org/en/stable/tutorials/notebooks/atac/PeakVI.html. The model reconstructed data is computed using the model.get_normalized_expression or model.get_normalized_accessibility function depending on the context.

#### multigrate

We conducted paired and trimodal integration using multigrate [25] following the tutorial listed in https://multigrate.readthedocs.io/en/latest/tutorials.html. To translate from modality 1 to modality 2, we obtained the latent representation of the observed modality using the model.module.encoder_0() function and then supplied it to the decoder of the unseen modality, model.module.decoder_1().

#### Evaluation of different methods

By default, we used binarized scATAC-seq data for multigrate, multiVI, peakVI, and MBTA. For scRNA-seq data, we used log-transformed, library size-normalized counts for multigrate and MBTA, and raw counts for scVI and multiVI. These differences are noted during the benchmarking process. The tested methods were evaluated in multiple ways. First, we computed the mean squared error between the observed, *o*, and the reconstructed/translated data, *o*^, for the observed modality for all samples: *MSE* = Σ*_i_*(*o_i_ − o*^*_i_*)^2^. In addition to sample-level statistics, we computed population-level statistics including the Cramér–von Mises (CvM) statistics, sample-sample correlation difference, and feature-feature correlation difference. When *O* and *O*^^^ are the full observed and reconstructed/translated data matrices, the CvM statistics are computed across features using the scipy.stats.cramervonmises_2samp() function.

The sample-sample correlation was computed between the rows of *O*and *O*^^^, yielding two correlation matrices, *R_s_* and R^*_s_*. To compute the sample-sample correlation difference, we substracted R^*_s_* from *R_s_*, and took its absolute value. The same procedure was used to compute the feature-feature correlation difference after computing the feature correlation between columns of *O* and *O*^^^. In both instances, we computed correlation matrices using the numpy.corrcoef() function on log normalized data.

To evaluate the quality of cell embeddings produced by each integration method, we used the bio-conservation metrics implemented in the scib-metrics package [74, 75].

### Constructing simulated datasets with scMultSim

Simulated co-profiled gene expression and chromatin accessibility data was constructed using the scMultSim package [26]. We simulated 1000 cells with 500 genes with either a tree-like or cluster-like topology. The tree topology was simulated by setting tree=Phyla5(), and the clustered data simulated by setting tree=ape::read.tree(text = “(A:1, B:1, C:1, D:1, E:1);”). Other parameters of the package were kept at their default values.

### Quantifying neighborhood structure similarities across datasets

Given two modalities, we constructed a k-nearest neighbor graph using each modality as input and subsequently quantified the similarity of the two KNN graphs by counting the number of shared edges. When the number of nodes (samples) was too large, we obtained a subsample from the data and constructed KNN graphs from smaller samples. When subsamples were taken, we repeated the process 100 times to ensure the robustness of our results.

### Inferring copy number alteration

InferCNV (https://github.com/broadinstitute/inferCNV) was used to perform CNV analysis on the scRNA-seq data that was aligned with the DNA-seq data. A raw gene-count matrix was generated from the aligned scRNA-seq data. Cells were annotated into reference and observation groups via the annotation file. Genes were ordered by genomic coordinates using a standard gene-position file (https://data.broadinstitute.org/Trinity/CTAT/cnv/). The cutoff was set to 0 to include all raw count scRNA-seq data. Discrete CNV state calling via the HMM was not activated.

CopyKat [41] was downloaded from https://github.com/navinlabcode/copykat and implemented following the package tutorial with default parameters.

### Quantifying the magnitude of cross-modal couplings

To quantify the magnitude of cross-modal coupling, we devised a perturbation-based approach. For each cell, we applied a random perturbation to its latent coordinate in modality 1. The magnitude of the perturbation was calibrated such that each latent dimension was displaced by 1 on average — a scale chosen to ensure the perturbation was large enough to propagate meaningfully through the decoder and produced a detectable change in the original feature space. Two parallel consequences of this perturbation were then quantified. First, both the original and the perturbed modality 1 latent coordinates were decoded back into the modality 1 feature space, and the resulting change was measured as the Euclidean distance between the two decoded outputs — providing the effective magnitude of the perturbation as expressed in modality 1. Second, both the original and the perturbed modality 1 latent coordinates were translated into the modality 2 latent space via MBTA and decoded into the modality 2 feature space, and the resulting change was again measured as a Euclidean distance — providing a quantification of the extent of the perturbation in modality 1 that propagates into modality 2. By comparing these two Euclidean distances, we obtained a direct readout of how strongly a change in modality 1 was reflected in modality 2. A large change in modality 2 relative to modality 1 indicates tight cross-modal coupling; a small change indicates that the two modalities respond largely independently. This analysis was performed per cell and can be aggregated across cell types or developmental stages to characterize the extent of variation of cross-modal coupling across the biological system.

### Constructing unimodal flows

MIOFlow [60] and TIGON [61] were used to construct unimodal flows that evolve a single modality. Both algorithms were downloaded from their publicly available online repositories (MIOFlow: https://github.com/KrishnaswamyLab/MIOFlow/tree/main/MIOFlow. TIGON: https://github.com/yutongo/TIGON) and trained following their respective tutorials with default parameters. Both algorithms were trained with modality-specific embeddings learned by MBTA.

### Construction of a flow map for embryonic development

Flow maps, Φ_Δ_*_t_*(*·*), allow the construction of a time-independent vector field that connects samples in time [59]. Given data points and their associated time coordinate, flow maps connect samples in time by optimal transport [76, 77]. Specifically, we conducted optimal transport on the latent space, *z*. Given data points from *z* and their associated time stamps, *{t_i_}*, we can represent the sample in the form of data matrices, *{Z_ti_ }*. A flow map is then constructed by minimizing the sliced Wasserstein distance between *Z_t_*_1_ and Φ*_t_*_2_ *_−t_*_1_ (*Z_t_*_1_). The sliced Wasserstein distance between two probability distributions, *p* and *q*, is defined as:

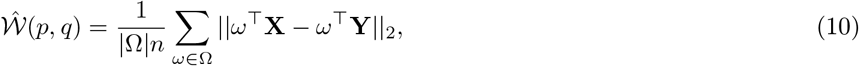

where *p* and *q* denote the probability distributions from which *X* and *Y* are sampled (assuming *ω^⊤^***X** and *ω^⊤^***Y** are ranked in ascending order), *n* is the dimension of the system, and Ω is a set of random projection directions.

### Introducing synthetic batch effects to data and batch correction with scVI

Batch effects were introduced to cells through the addition of a gene-specific offset. For each gene, we first computed its mean expression in cells to which a batch effect was to be introduced. Then, we selected a random number for each gene between -1 to 5 via np.random.randint; this number is termed the gene-specific multiplier. Finally, we multiplied the average expression of the gene by the gene-specific multiplier and added this gene-specific offset to each cell. Any number less than 0 after the addition was set to 0 and the final result was rounded to generate count data.

Single-cell RNA-seq data were integrated using scVI to remove batch effects across simulated batches. The scVI model was configured with 4 encoder/decoder layers, a latent dimensionality of 30, hidden layer width of 600, and a negative binomial gene counts likelihood. The model was trained for a maximum of 1,000 epochs on a single GPU. The resulting batch-corrected gene expression was obtained by model.get_normalized_expression(transform_batch=’reference’) and was used as input for MBTA.

### Application of single cell foundation models

Geneformer [62] and UCE [64] were implemented via the omicverse [78] platform with default settings. scGPT [63] was installed from https://github.com/bowang-lab/scGPT and embeddings were generated via the command scg.tasks.embed_data. STATE [65] was first installed from https://github.com/ArcInstitute/state and the model check point was downloaded from https://huggingface.co/arcinstitute/SE-600M. The STATE embedding was generated via the command uv run state emb fit.

### Data preprocessing

Gene expression data counts were normalized by library size and subsequently log-transformed using the sc.pp.normalize_total and sc.pp.log1p function in scanpy [79]. Chromatin accessibility and epigenetic profiles were binarized. Protein abundance values from CITE-seq experiments were z-scored. Copy number profiles from the ER+ breast tumor samples were in the consensus segment ratio format and required no additional preprocessing.

### Data availability

All datasets used in this paper are publicly available. The PBMC dataset was downloaded from the multiVI tutorial page at https://docs.scvi-tools.org/en/stable/tutorials/notebooks/multimodal/MultiVI_tutorial.html. The BMCC dataset was downloaded from the multigrate tutorial page at https://multigrate.readthedocs.io/en/latest/notebooks/paired_integration_cite-seq.html. The hybrid cell and mouse brain dataset [18] were downloaded from https://ngdc.cncb.ac.cn/bioproject/browse/PRJCA024774. The HSPC dataset was downloaded from https://www.kaggle.com/competitions/open-problems-multimodal/data. The developing embryo dataset [19] was downloaded from https://www.ncbi.nlm.nih.gov/geo/query/acc.cgi?acc=GSE235109 and https://www.ncbi.nlm.nih.gov/geo/query/acc.cgi?acc=GSE259393. The ER+ breast cancer dataset [17] can be downloaded from https://www.ncbi.nlm.nih.gov/sra/?term=PRJNA1086561 and https://www.ncbi.nlm.nih.gov/geo/query/acc.cgi?acc=GSE261713.

## Code availability

Code used for this project has been deposited on Github (https://github.com/bxxu/move_between_modalities).

