## supplemental figures and notes for "Move BeTween modAlities (MBTA) employs flow matching to predict single cell data modalities"

### Supplementary note 1: Flow matching

The flow matching component of MBTA constructs flow fields between any arbitrary choice of two modalities with latent spaces z1 and z2. The expected coordinate of a cell, $y$, on $z_2$ given its coordinate on $z_1$, $x$, is the solution of the ordinary differential equation, $\frac{dy}{dt} = f_{z_1 \to z_2}(x, y, t)$.

#### Intermediate state in the flow path

A potential source of confusion in interpreting the flow matching component of MBTA is the role of the intermediate states generated. Because flow matching is parameterized as a neural ordinary differential equation (ODE), it produces a continuous path from an initial distribution to a target distribution. This path passes through the latent space, and it is natural to ask whether these states carry biological meaning — for example, whether they represent intermediate cell states, developmental transitions, or regulatory intermediates between modality 1 and modality 2. They do not. The intermediate states are purely auxiliary mathematical constructs that arise from the numerical integration of the ODE and have no biological interpretation. The only value that matters is $y(t = 1)$, which is the generated coordinate in the space of $z_2$ given a $z_1$ coordinate $x$.

#### Dealing with one-to-many biological relationships

The flow matching component of MBTA learns the conditional distribution ($p(z_2 \mid z_1)$) and generate samples in the space of $z_2$ using $z_1$ as conditional input via integration along the flow path defined by $\frac{dy}{dt} = f_{z_1 \to z_2}(x, y, t)$. As a result, MBTA should be interpreted as generating the probability distribution of modality 2 states conditioned on modality 1. The contrast of flow matching with regression based approach is intuitive. A regression model, $f_R$, minimizes $\mathbb{E}[|z_2 - f_R(z_1)|^2]$, whose solution is the conditional mean $f_R(z_1) = \mathbb{E}[z_2 \mid z_1]$ — a single point that averages over all possible $z_2$ values consistent with a given $z_1$. When the conditional distribution $p(z_2 \mid z_1)$ is multimodal, this average falls between the modes and corresponds to no real state in the data. Flow matching, by contrast, transports probability mass toward the full conditional distribution $p(z_2 \mid z_1)$. All modes are preserved: a given $z_1$ gives rise to a distribution over $z_2$ values rather than a single point estimate, and distinct high-probability regions in the target latent space remain distinct after transport. When the relationship between different molecular modalities is not strictly one-to-one, this formulation can preserve the heterogeneity of data.

To support this point, we extensively benchmarked MBTA using co-profiled datasets. These datasets enable direct evaluation of cross-modal translation accuracy because both modalities are measured in the same single cells. Specifically, when MBTA translates modality 1 into modality 2, we compute the mean squared error (MSE) between the translated modality 2 and the true modality 2 measured from that same cell. Across datasets, MBTA consistently achieved the strongest translation performance, with lower MSE, than competing methods while also better preserving the original cell–cell and feature-feature relationships, as quantified by cell–cell and feature-feature correlation.

#### Conditional flow matching

MBTA essentially translates between modalities via conditional flow matching. A natural alternative design would be to initialize the flow directly from $z_1$ rather than from Gaussian noise, using the source latent coordinate as the starting point of the trajectory toward $z_2$. This would in principle allow the flow to exploit the geometric structure already encoded in $z_1$, potentially shortening the path to $z_2$ and improving efficiency. This approach also carry an appealing theoretical property: because the source information is encoded in the starting position rather than as a conditioning signal, the vector field itself becomes cell-independent — a single, shared function of position and time that describes all inter-modality translations, rather than a cell-specific field that must be re-evaluated for each new input. However, this design is only feasible when $z_1$ and $z_2$ share the same dimensionality — a restriction that would severely limit the generality of the framework. In MBTA, the two VAEs are trained independently, and the dimensionality of each latent space is chosen to reflect the complexity of its own modality: a compact space may suffice for a low-dimensional protein readout, while a higher-dimensional space may be warranted for a genome-wide chromatin accessibility profile. When $z_1$ and $z_2$ have different dimensions, using $z_1$ as an initial condition for a flow in $z_2$ space is geometrically undefined. The Gaussian initialization resolves this by providing a universal, dimension-agnostic starting distribution, with $z_1$ entering the model as a conditional input that guides the trajectory rather than as its origin. This is not a simplification but a deliberate design choice that preserves the full generality of the framework.

#### Supplementary note 2: Computational efficiency and choice of hyperparameters

The training of MBTA consists of two steps: training of the VAE and training of the flow matching components. In general, we observed that larger datasets typically take fewer epochs to train, as was noted earlier [30], and that the training of the VAE component largely depends on the dimensionality of the data. To analyze the efficiency of MBTA training, we generated synthetic co-profiled datasets with a feature space that spans between 20 to 100,000 genes/peaks and trained 5 separate models for each synthetic dataset. All training was conducted on a server equipped with dual Intel Xeon E5-2683 v3 processors. Training of neither component required parallel computation. Two molecular modalities (for instance, scRNA-seq data of 1000 genes and scATAC-seq data containing 10000 peaks) need two VAE components and two flow matching components, taking roughly an hour to train (Figure A1A, B).

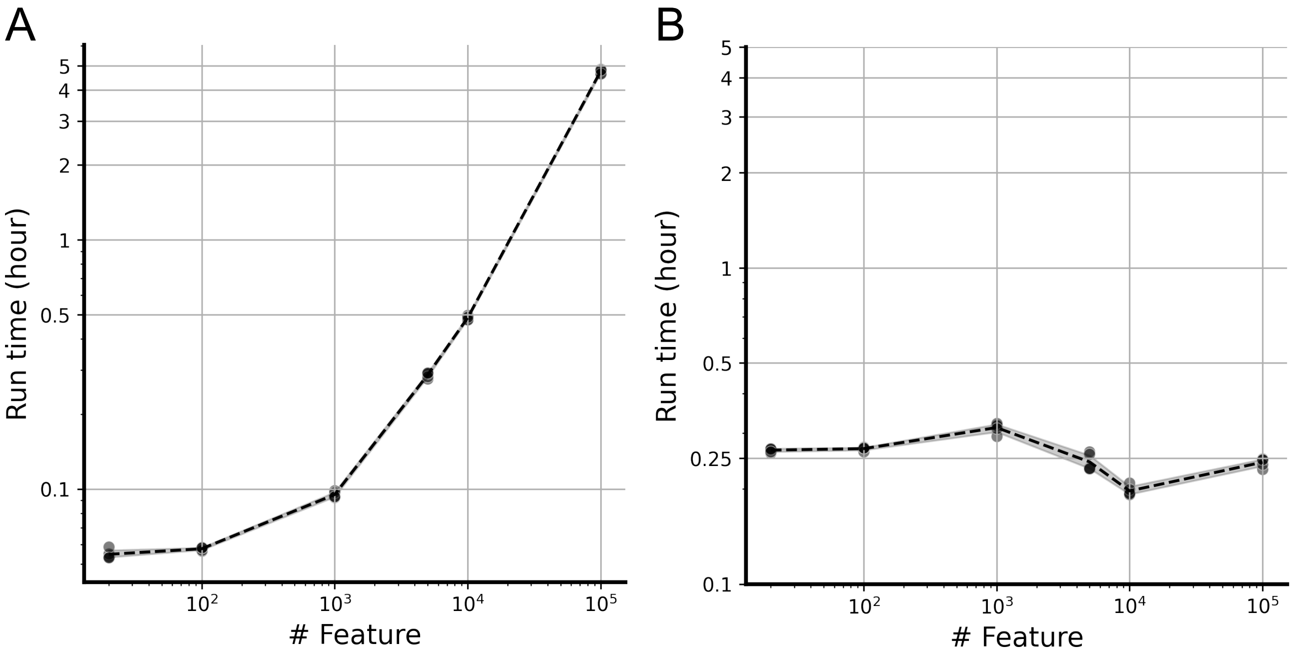


Figure A1: Training time of the VAE component (A) and the flow matching component (B)

The quality and topology-preserving properties of the VAE latent space are governed by two key architectural parameters: the latent space dimensionality and the neural network width. The latent dimensionality determines the degrees of freedom available to represent cellular heterogeneity — a space that is too low-dimensional will compress distinct cell states into overlapping representations, while one that is too high-dimensional may overfit to noise and produce a poorly regularized latent geometry. The neural network width, on the other hand, controls the expressivity of the encoder and decoder: wider networks can capture more complex, nonlinear relationships between the input feature space and the latent space, but at the risk of overfitting when the network capacity greatly exceeds what the data requires.

To evaluate how sensitive the downstream flow matching performance is to these choices, we trained MBTA models across a range of hyperparameter combinations, systematically varying both the latent space dimensionality and the neural network width of the VAE components. For each combination, we repeated training five times to separate the effect of hyperparameter choice from stochastic variation. Through this approach, we found that the number of neural network nodes has a greater impact on model performance than the number of latent dimensions (Figure A2–A5). When the number of nodes is in the range of 100–200, performance is largely insensitive to the exact choice of latent dimensionality (Figure A2–A5). Notably, using too many neural network nodes increases MSE, likely due to overfitting. These results suggest that MBTA is robust to reasonable choices of VAE architecture, and that the flow matching component is not critically sensitive to the precise geometry of the latent space, provided that the VAE is not severely under- or over-parameterized.

To balance this opposing trend, we use a 20 dimensional latent space with 200 nodes on each neural network layer by default (Figure A2–A5).

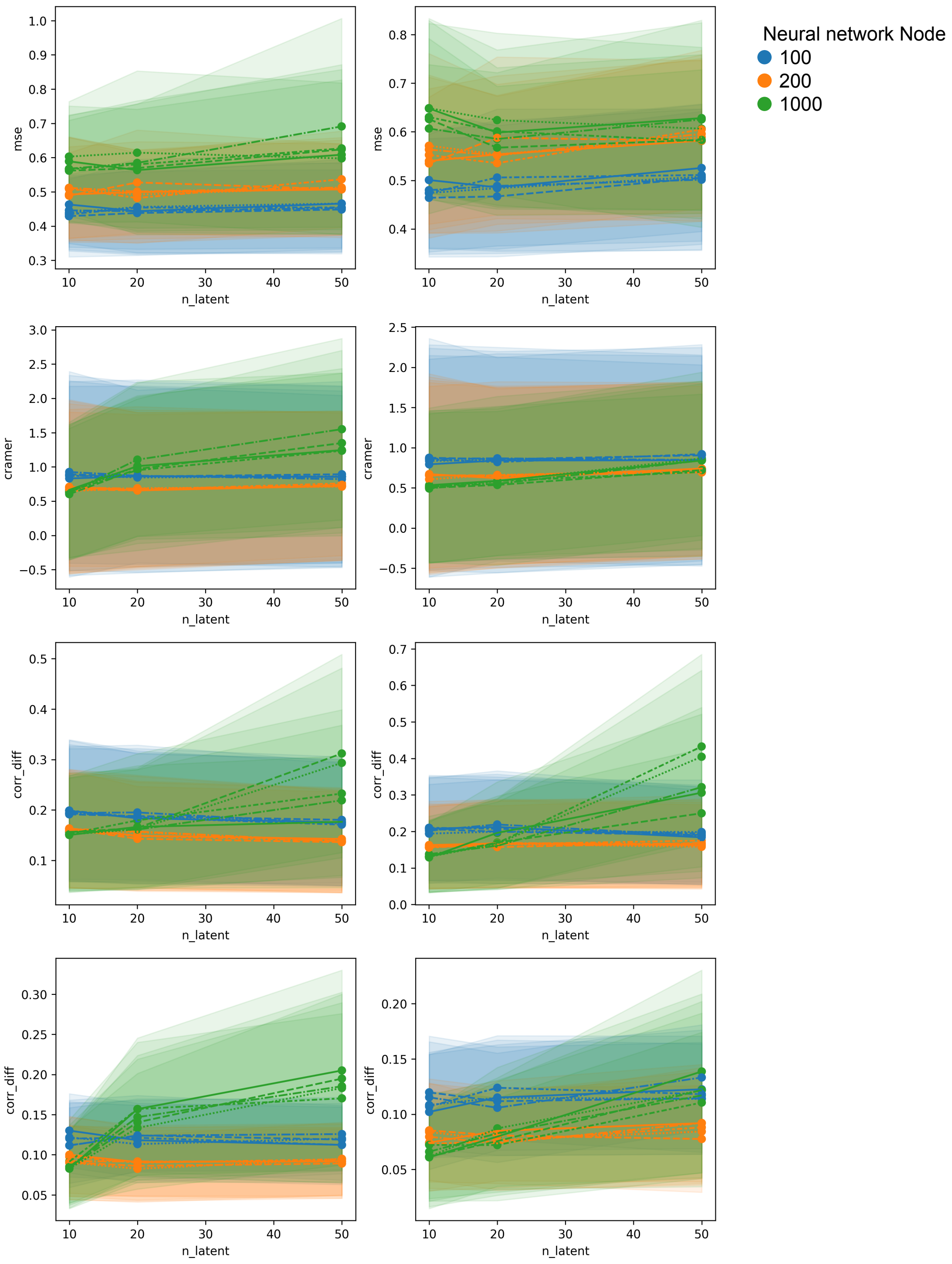


Figure A2: The line plots illustrate the mean squared error (MSE), CvM statistics, sample correlation difference and feature correlation difference for data reconstruction (left column) and data translation (right column) using MBTA models trained on the synthetic cluster dataset. The x-axis represents the latent space dimension, ranging from 10 to 50. Colors denote the neural network width, with blue, orange, and green representing 100, 200, and 1000 nodes, respectively. Each dot corresponds to a distinct model trained under specific initializations, and the shaded regions represent the variability of all the test cells. Line style denotes replicate model.

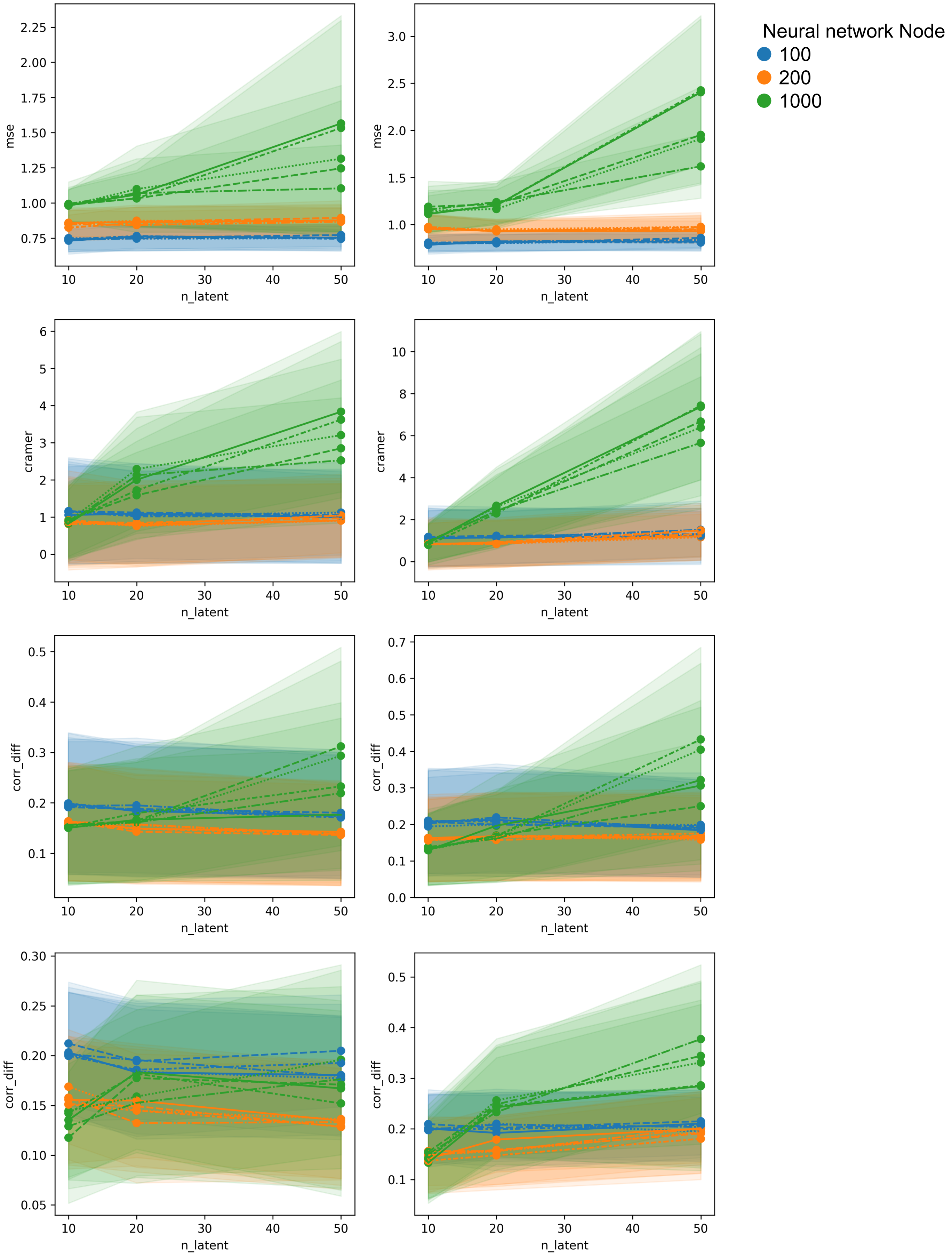


Figure A3: The MSE, CvM, feature correlation difference and sample correlation difference of the synthetic tree dataset. Line style denotes replicate model.

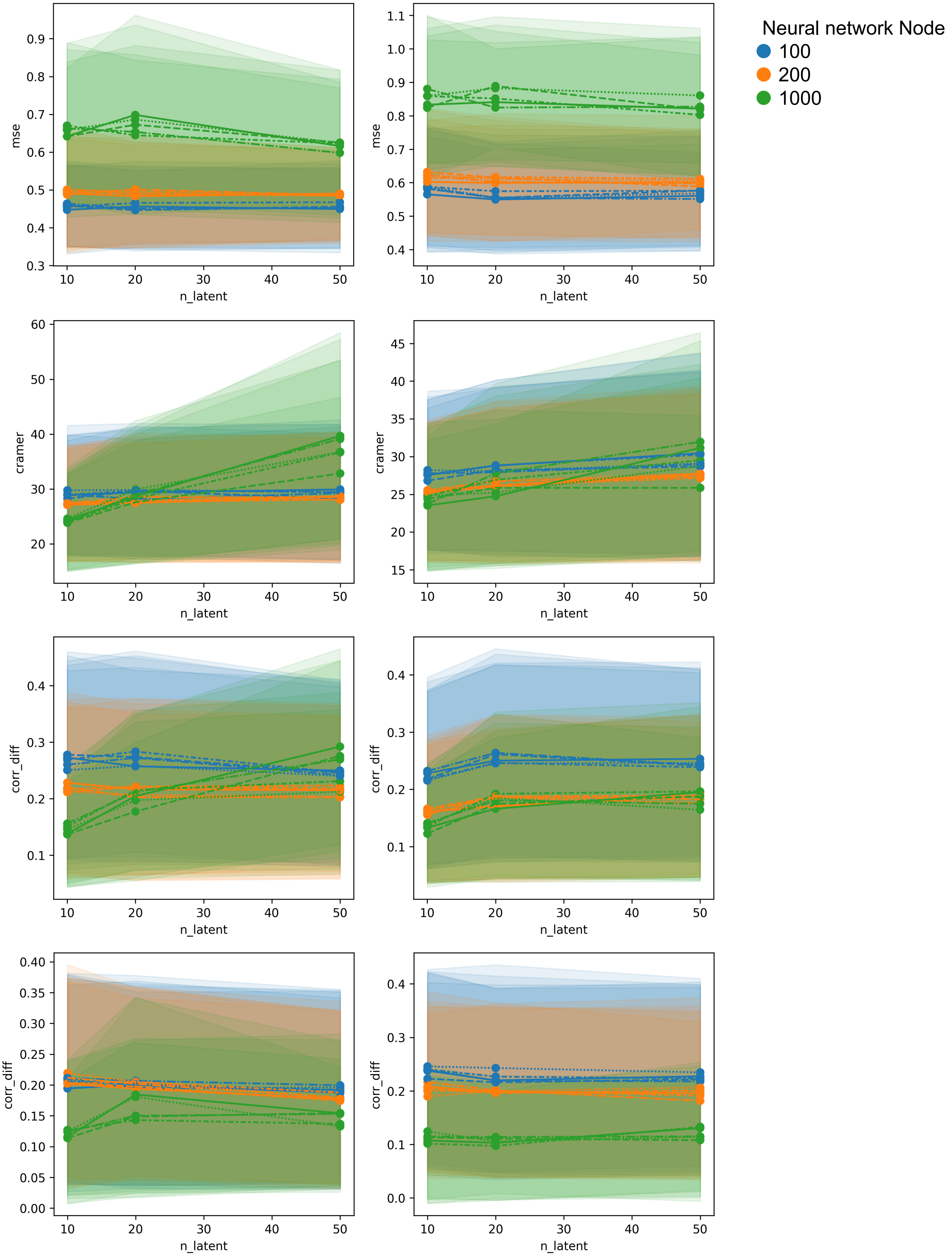


Figure A4: The MSE, CvM, feature correlation difference and sample correlation difference of the hybrid cell dataset. Line style denotes replicate model.

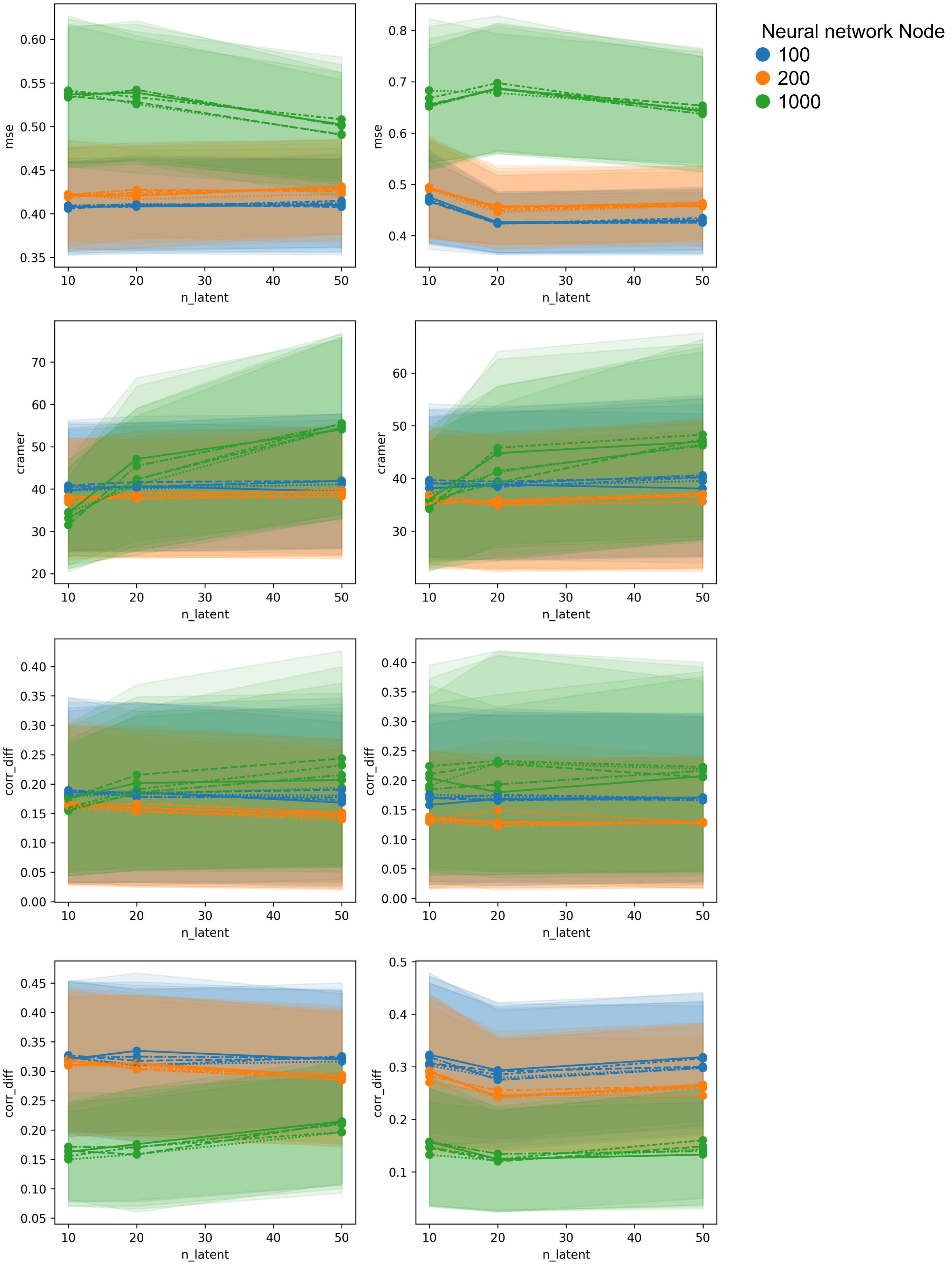


Figure A5: The MSE, CvM, feature correlation difference and sample correlation difference of the PBMC dataset. Line style denotes replicate model.

#### Supplementary note 3: Using foundation models to address structural mismatch

A natural question is whether the structural mismatch problem that MBTA is designed to address might already be resolved by foundation models trained on large-scale single-cell data. Foundation models leverage extensive data coverage and high representation capacity, and it is reasonable to ask whether scale alone might mitigate modality-specific heterogeneity — rendering explicit modeling choices such as those in MBTA unnecessary.

To address this question, we first consider the training objectives of current foundation models for single-cell genomics, including models trained on gene expression (Geneformer [62], scGPT [63], UCE [64], STATE [65]) and chromatin accessibility (EpiFoundation [80], EpiAgent [81], ChromFound [82]). These models have demonstrated strong performance in large-scale representation learning and in downstream tasks such as cell type clustering, annotation, and retrieval. However, their primary training objectives — masked feature prediction, contrastive learning, and embedding alignment — are not designed to support end-to-end reconstruction or cross-modal translation in the original feature space.

In masked prediction, a proportion of genes or genomic regions in the input are randomly hidden, and the model is trained to predict the missing values from the remaining context. This teaches the model to capture dependencies between features but does not enable decoding of the full cell profile from the learned embedding. In contrastive learning and embedding alignment, the model is trained to bring representations of related samples — for example, the RNA and ATAC profiles of the same cell — closer together in embedding space while pushing unrelated samples apart. This produces well-aligned embeddings across modalities, but the model acquires no capacity to decode those embeddings back into the original feature space. As a result, none of these training schemes enables the reconstruction or modality translation tasks that MBTA is designed to address.

A related but architecturally distinct class of models is represented by AlphaGenome [83], which takes genomic DNA sequence as input and predicts multiple molecular readouts — including gene expression, chromatin accessibility, histone modifications, and transcription factor binding — through modality-specific prediction heads. AlphaGenome operates strictly in the sequence-to-modality direction: given a DNA sequence, it predicts the molecular landscape of the cell. This is a powerful capability for understanding the regulatory grammar encoded in the genome, but it is architecturally incompatible with the cross-modal translation task addressed here. AlphaGenome cannot take an observed RNA profile and predict the corresponding ATAC profile, nor can it take an observed ATAC profile and predict the corresponding RNA profile.

MBTA is specifically designed for reconstruction and cross-modal translation, with explicit modeling choices aimed at handling structural mismatch — the phenomenon in which the neighborhood structure of a cell differs substantially across modalities. Most current foundation models are optimized to learn informative latent representations, and additional decoder components would need to be introduced to map those embeddings back to the original data space for reconstruction or translation. While such extensions are possible, a recent benchmarking study [84] reported that simple reconstruction strategies from foundation-model embeddings underperformed even PCA-based baselines, suggesting that strong embeddings do not automatically translate into strong generative performance and that substantial additional development would be required to achieve competitive reconstruction accuracy.

More broadly, the premise that scale alone might resolve the limitations of current foundation models has been directly challenged by a recent systematic study [85]. DenAdel et al. pretrained 400 single-cell foundation models across a corpus of 22.2 million cells and evaluated performance across zero-shot cell type classification, batch integration, and perturbation response prediction. They found that performance plateaus at a small fraction of the available training data, and that — unlike large language models — single-cell foundation models exhibit no clear data scaling laws. These results indicate that the limitations of current foundation models in single-cell biology reflect fundamental constraints that are not resolved by increasing training data volume.

To assess empirically whether large pretrained unimodal foundation models might resolve structural mismatch through scale alone, we evaluated STATE, UCE, scGPT, and Geneformer on three datasets commonly used to benchmark multimodal integration methods. Across all three datasets, the resulting embeddings did not preserve biologically meaningful heterogeneity as effectively as expected, scoring on average 4.4% worse than generic gene expression UMAPs in cell type annotation tasks (Figure A6–A9). These results indicate that increased data scale and representation capacity alone do not resolve the structural mismatch problem.

Taken together, these analyses suggest that current foundation models and MBTA serve related but distinct purposes. Foundation models are highly valuable for large-scale representation learning, but scale alone does not fully address cross-modal structural mismatch. This supports the need for explicit modeling strategies — such as those implemented in MBTA — for accurate reconstruction and modality translation.

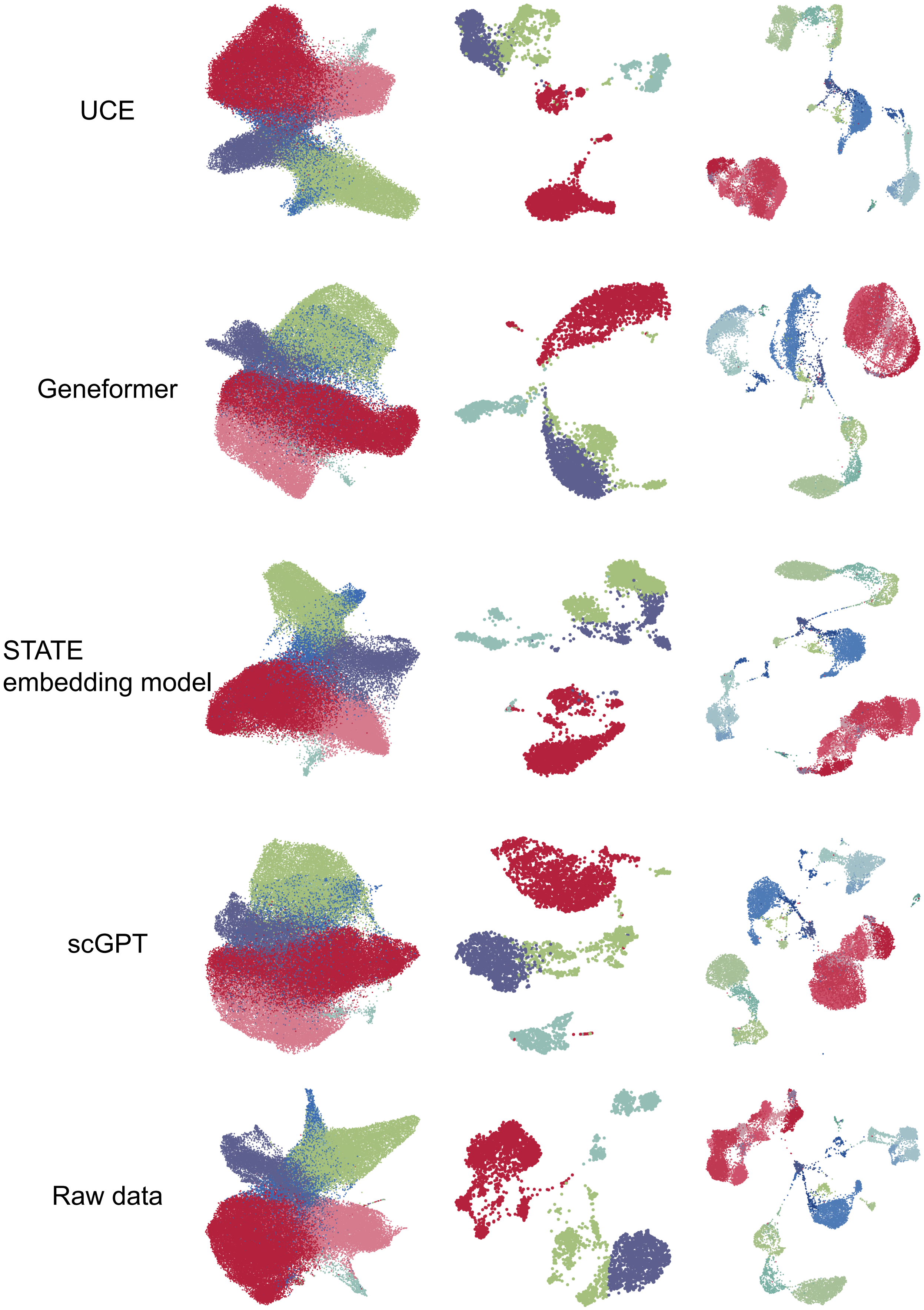


Figure A6: UMAP of embeddings generated by multiple foundation models on the HSPC (first column), PBMC (second column) and BMCC (third column) datastes. Their respective embedding qualities are summarized in figure A7, A8 and A9 respectively.

#### HSPC

| Method | Bio conservation: Isolated labels | Bio conservation: KMeans NMI | Bio conservation: KMeans ARI | Bio conservation: Silhouette label | Bio conservation: cLISI | Aggregate score: Bio conservation |
|---|---|---|---|---|---|---|
| **STATE umap** | 0.55 | 0.56 | 0.41 | 0.60 | 0.99 | 0.62 |
| **RNA umap** | 0.56 | 0.54 | 0.38 | 0.59 | 0.98 | 0.61 |
| **STATE** | 0.51 | 0.56 | 0.40 | 0.52 | 0.98 | 0.59 |
| **UCE umap** | 0.53 | 0.49 | 0.36 | 0.58 | 0.97 | 0.58 |
| **UCE** | 0.52 | 0.49 | 0.36 | 0.53 | 0.97 | 0.57 |
| **scGPT** | 0.52 | 0.45 | 0.35 | 0.53 | 0.99 | 0.57 |
| **scGPT umap** | 0.53 | 0.45 | 0.33 | 0.54 | 0.95 | 0.56 |
| **Geneformer umap** | 0.52 | 0.46 | 0.32 | 0.54 | 0.95 | 0.56 |
| **Geneformer** | 0.52 | 0.44 | 0.32 | 0.52 | 0.97 | 0.55 |

Figure A7: Embedding quality of gene expression and those generated by different foundation models for the HSPC dataset. Biological conservation metrics were computed on the embeddings of each method. "Isolated labels" quantifies whether each annotated cell type is distinct from all other cell types. "KMeans NMI" and "KMeans ARI" measure the agreement between k-means clusters derived from the embedding and known cell type annotations, using normalized mutual information and the adjusted Rand index respectively. "Silhouette label" measures the compactness and separation of cell type clusters in the embedding. "cLISI" (cell-type Local Inverse Simpson Index) quantifies the purity of each cell's local neighborhood with respect to cell type identity, with a score of 1 indicating a perfectly pure neighborhood. For all columns, a higher score indicates better embedding quality. Aggregate score is computed as the average of the five scores.

#### PBMC

| Method | Bio conservation: Isolated labels | Bio conservation: KMeans NMI | Bio conservation: KMeans ARI | Bio conservation: Silhouette label | Bio conservation: cLISI | Aggregate score: Bio conservation |
|---|---|---|---|---|---|---|
| **Geneformer umap** | 0.75 | 0.77 | 0.81 | 0.75 | 1.00 | 0.81 |
| **scGPT** | 0.62 | 0.83 | 0.90 | 0.62 | 1.00 | 0.79 |
| **UCE umap** | 0.78 | 0.70 | 0.64 | 0.79 | 1.00 | 0.78 |
| **scGPT umap** | 0.81 | 0.73 | 0.55 | 0.80 | 1.00 | 0.78 |
| **RNA umap** | 0.78 | 0.72 | 0.59 | 0.79 | 1.00 | 0.78 |
| **STATE umap** | 0.79 | 0.74 | 0.54 | 0.79 | 1.00 | 0.77 |
| **STATE** | 0.54 | 0.80 | 0.86 | 0.54 | 1.00 | 0.75 |
| **UCE** | 0.58 | 0.70 | 0.59 | 0.57 | 1.00 | 0.69 |
| **Geneformer** | 0.54 | 0.57 | 0.49 | 0.54 | 1.00 | 0.63 |

Figure A8: Embedding quality of gene expression and those generated by different foundation models for the PBMC dataset.

BMCC

| Method | Bio conservation: Isolated labels | Bio conservation: KMeans NMI | Bio conservation: KMeans ARI | Bio conservation: Silhouette label | Bio conservation: cLISI | Aggregate score: Bio conservation |
|---|---|---|---|---|---|---|
| **STATE umap** | 0.64 | 0.72 | 0.44 | 0.56 | 1.00 | 0.67 |
| **RNA umap** | 0.63 | 0.72 | 0.41 | 0.55 | 1.00 | 0.66 |
| **UCE** | 0.56 | 0.71 | 0.45 | 0.55 | 1.00 | 0.65 |
| **UCE umap** | 0.61 | 0.68 | 0.37 | 0.57 | 1.00 | 0.65 |
| **STATE** | 0.53 | 0.72 | 0.44 | 0.53 | 1.00 | 0.64 |
| **scGPT** | 0.56 | 0.68 | 0.42 | 0.55 | 1.00 | 0.64 |
| **scGPT umap** | 0.59 | 0.67 | 0.35 | 0.54 | 0.99 | 0.63 |
| **Geneformer** | 0.54 | 0.63 | 0.38 | 0.54 | 1.00 | 0.62 |
| **Geneformer umap** | 0.58 | 0.63 | 0.30 | 0.53 | 0.99 | 0.60 |

Figure A9: Embedding quality of gene expression and those generated by different foundation models for the BMCC dataset.

| Method | GitHub Repo | Input Format | Reconstruction capability |
|---|---|---|---|
| EpiFoundation [80] | https://github.com/UCSC-VLAA/EpiFoundation | binned chromatin accessibility | Reconstruct binarized gene expression |
| EpiAgent [81] | https://github.com/xy-chen16/EpiAgent | ranked cCRE accessibility | cCRE-space reconstruction |
| ChromFound [82] | https://github.com/SAIS-LifeScience/ChromFound | Binned ATAC-seq data + cell type label | embedding only |
| Geneformer [62] | https://huggingface.co/ctheodoris/Geneformer | Ranked gene tokens | embedding only |
| scGPT [63] | https://github.com/bowang-lab/scGPT | RNA-seq + gene tokens | embedding only |
| STATE [65] | https://github.com/ArcInstitute/state | RNA-seq | embedding only |
| UCE [64] | https://github.com/snap-stanford/UCE | RNA-seq | embedding only |
| SCARF [86] | https://github.com/cbmi-group/scarf | embedding only | No (requires reference RNA-seq data) |

Table A1: Summary table of foundation models, input formats, and prediction capabilities.

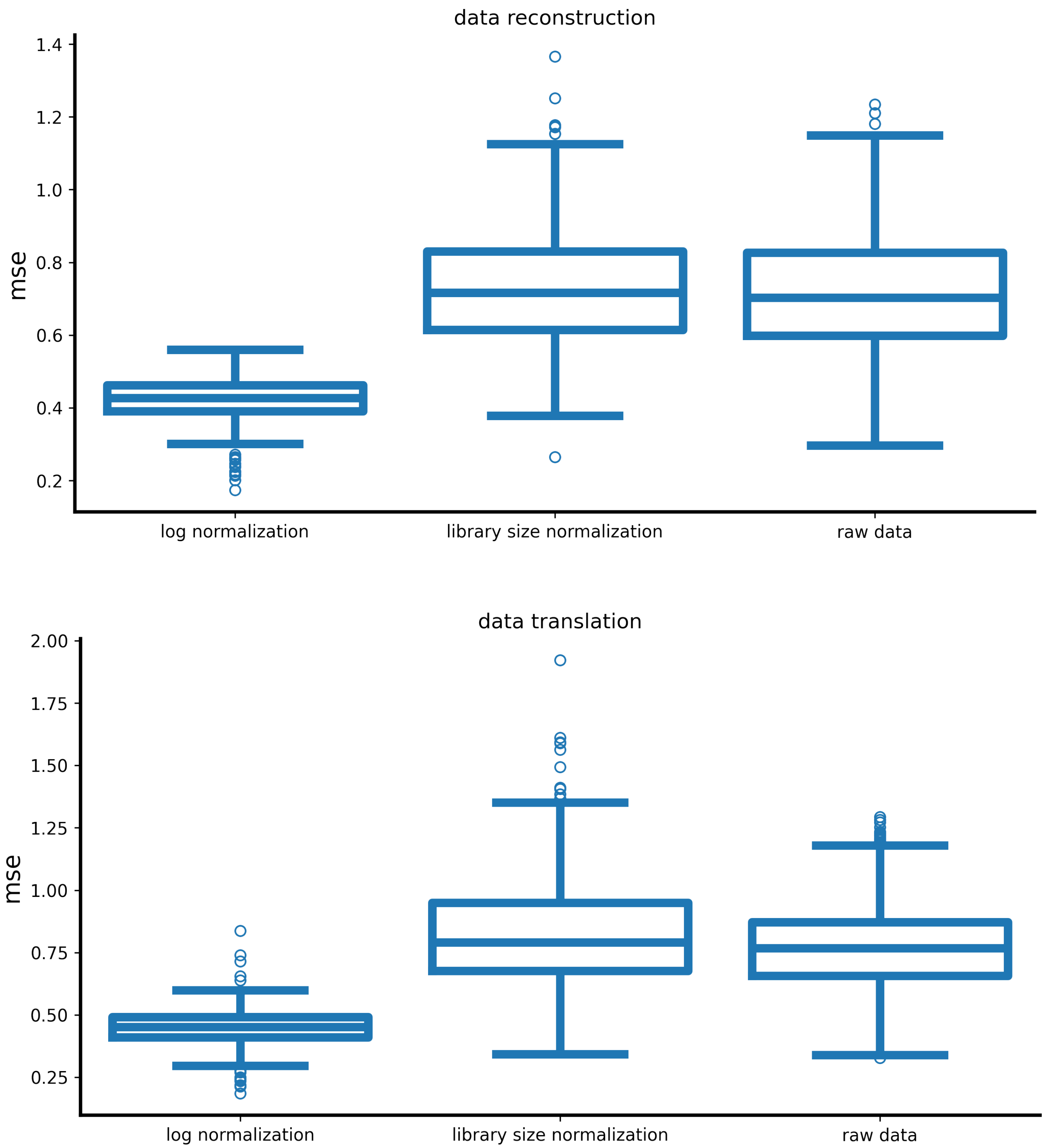


Figure S1: Data reconstruction and translation MSE for the PBMC dataset when the input data is preprocessed with different normalization approaches.

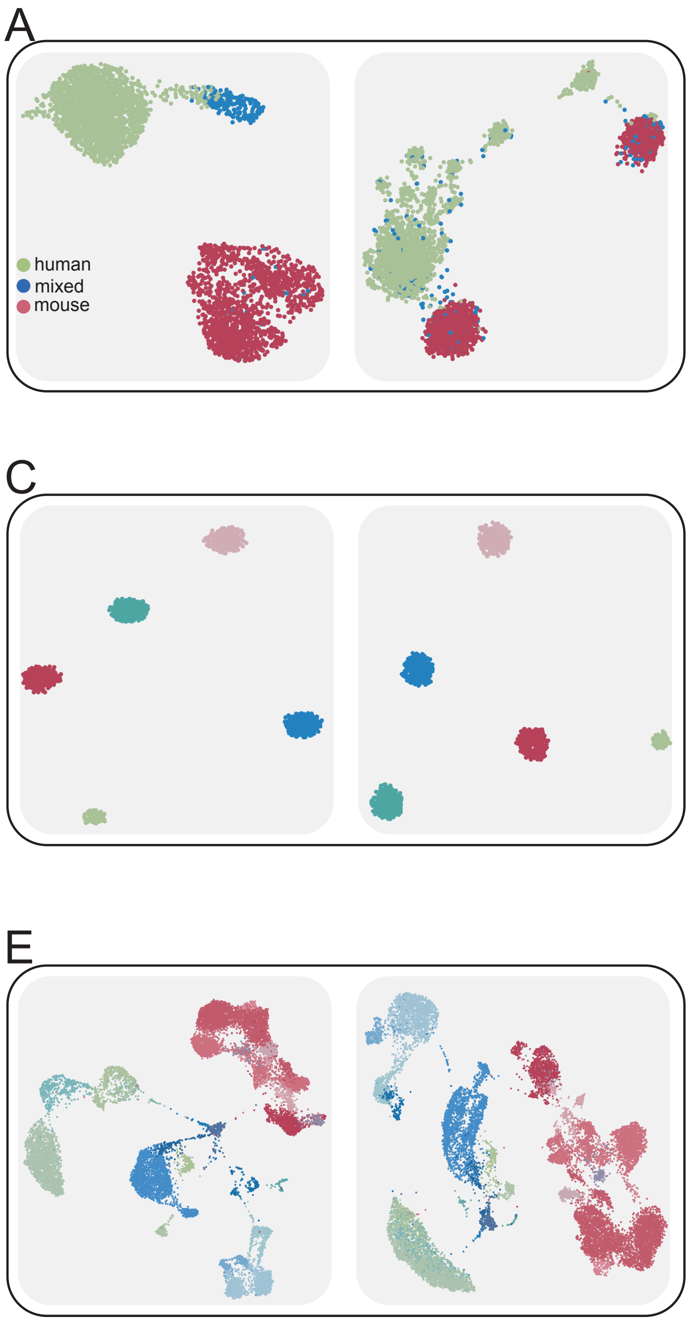


Figure S2: UMAP projection of the hybrid cell (A), PBMC (B), synthetic cluster (C), synthetic tree (D) and the BMCC (E) datasets. Colors denote cluster assignment, and each grid denotes one modality. All datasets used are co-profiled.

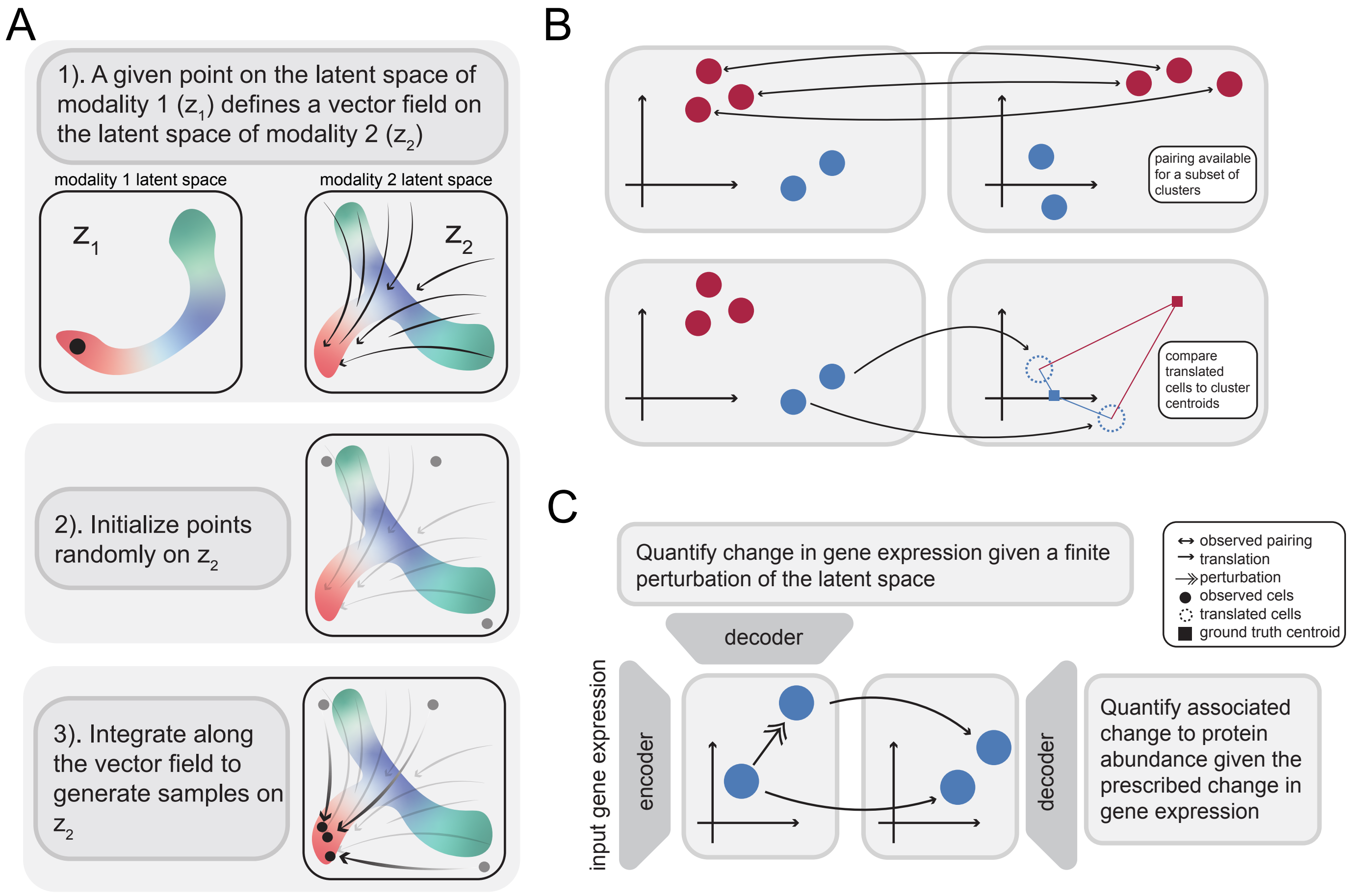


Figure S3: A: Detailed workflow of how MBTA uses the latent coordinate of one cell (shown in black) to translate it to the other modality. B: Illustration of partial pairing, where only a subset of the samples are co-profiled. C: Illustration of the analysis of modality coupling used to quantify how changes in one modality relate to changes in the other modality.

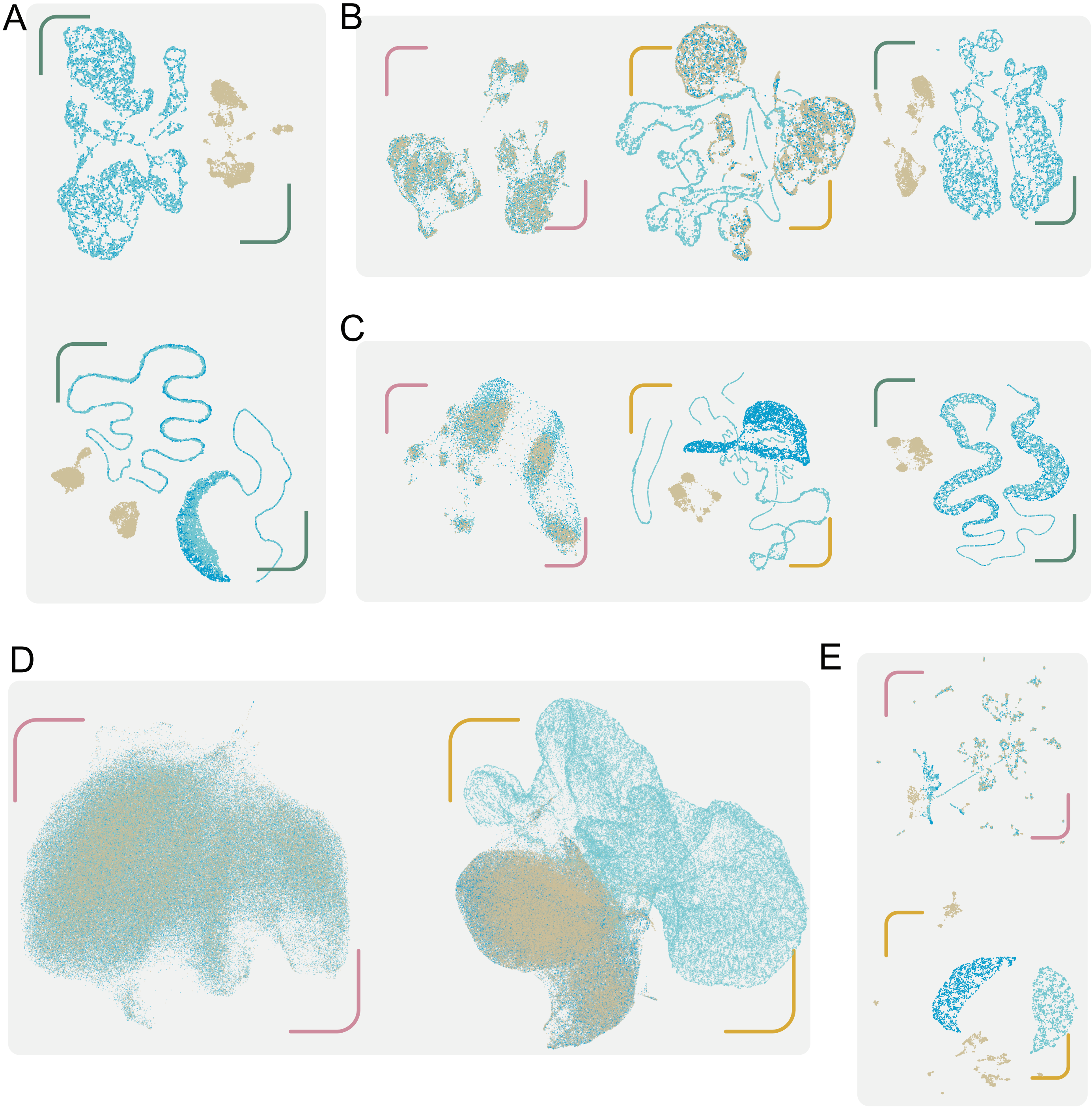


Figure S4: A: UMAP co-embedding of the hybrid cell and PBMC data constructed from multiVI. B: UMAP co-embedding of chromatin accessibility of the PBMC dataset constructed using MBTA, multigrate and multiVI, respectively. C: UMAP co-embedding of chromatin accessibility of the hybrid cell dataset constructed using MBTA, multigrate and multiVI, respectively. D: UMAP co-embedding of the protein abundance data from the HSPC dataset. E: UMAP co-embedding of the copy number data from the ER+ breast cancer data.

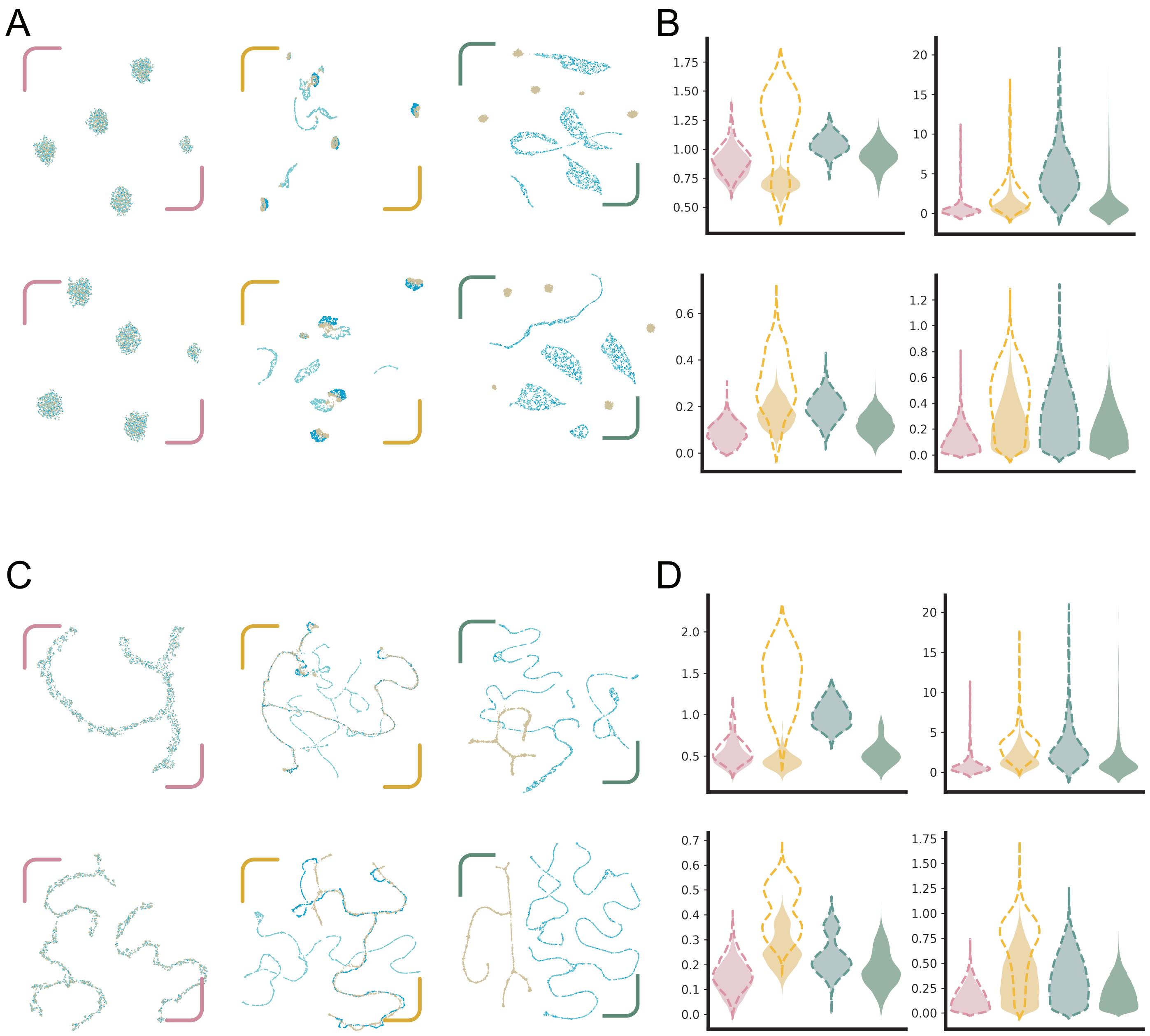


Figure S5: A: UMAP co-embeddings of both modalities from the synthetic cluster dataset using MBTA, multigrate and multiVI. B: Quantification of model performance using MSE, CvM, sample correlation difference and feature correlation difference. C: UMAP co-embeddings of both modalities from the synthetic tree dataset using MBTA, multigrate and multiVI. D: Quantification of model performance using MSE, CvM, sample correlation difference and feature correlation difference.

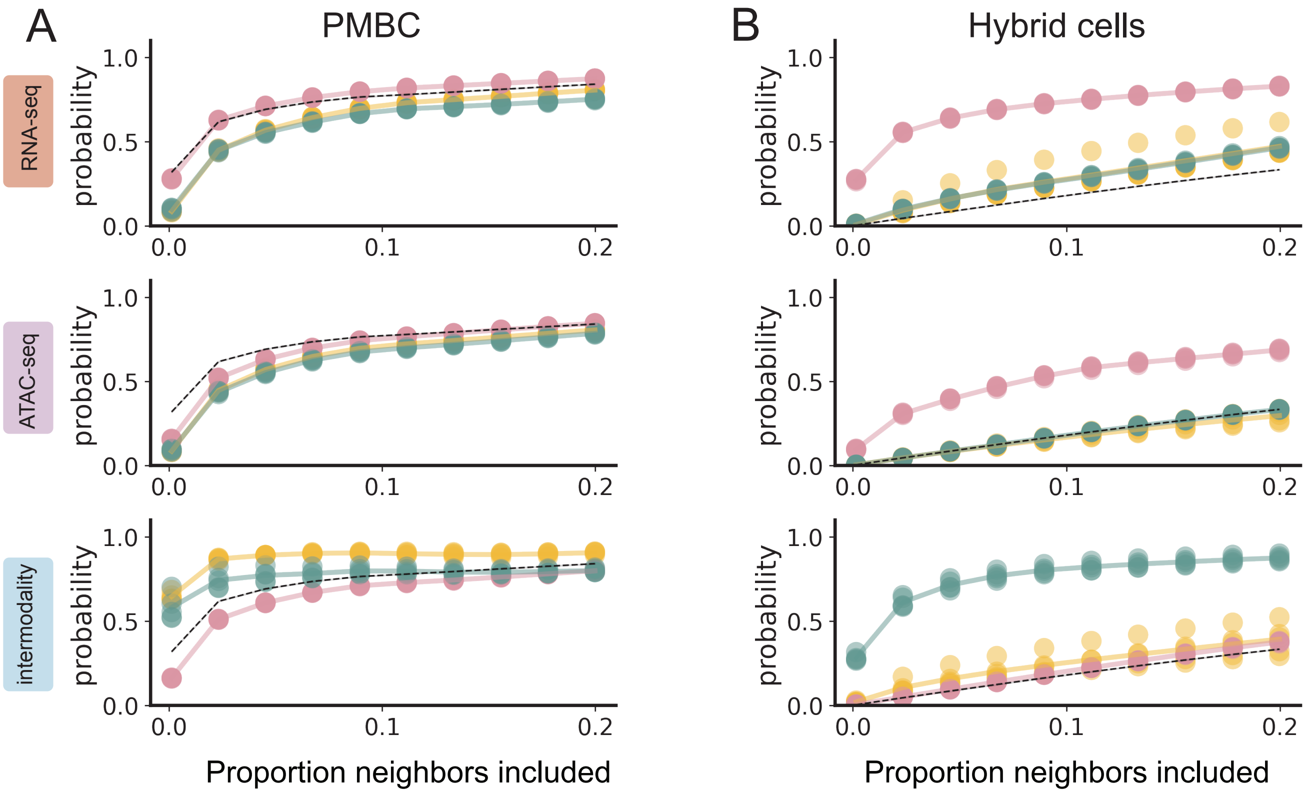


Figure S6: Similarity between KNN graphs constructed from the observed and reconstructed gene expression (first row), the observed and reconstructed chromatin accessibility (second row), and the reconstructed gene expression and the reconstructed chromatin accessibility (third row). The black dashed line represents the baseline intermodal structural mismatch computed from the observed gene expression and the observed chromatin accessibility. Results are computed from the PBMC (A) and hybrid cells (B) datasets.

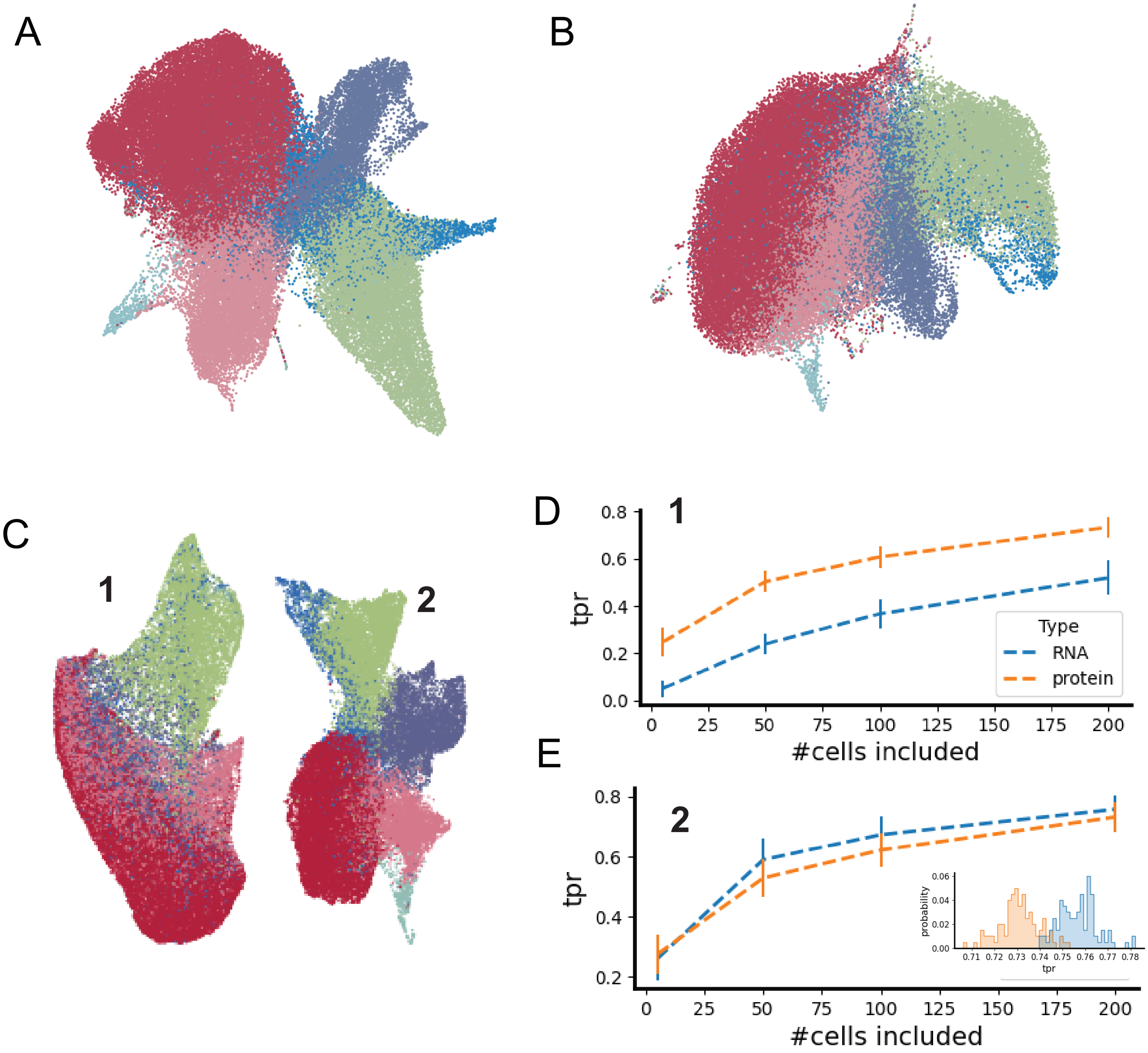


Figure S7: UMAP projection of the RNA space (A), protein space (B) and multigrate latent space (C) of the HSPC dataset. C-E, KNN graphs are constructed from the RNA data, protein data and multigrate latent space coordinates using either the cluster 1 or cluster 2 cells. For each cluster, both the RNA and the protein KNN graphs are compared against the latent space KNN graph (D,E). For cluster 1 cells, the protein graph is more similar to the latent space graph (D). For cluster 2 cells, the RNA graph is more similar to the latent space graph; the inset shows the distribution of actual true positive ratio (E).

A

| Method | Bio conservation: Isolated labels | Bio conservation: KMeans NMI | Bio conservation: KMeans ARI | Bio conservation: Silhouette label | Bio conservation: cLISI | Aggregate score: Bio conservation |
|---|---|---|---|---|---|---|
| **RNA umap** | 0.71 | 0.70 | 0.63 | 0.66 | 1.00 | 0.74 |
| **multiVI** | 0.70 | 0.67 | 0.59 | 0.68 | 1.00 | 0.73 |
| **MBTA umap** | 0.67 | 0.64 | 0.61 | 0.65 | 1.00 | 0.71 |
| **multiVI umap** | 0.67 | 0.64 | 0.53 | 0.65 | 1.00 | 0.70 |
| **MBTA** | 0.55 | 0.50 | 0.44 | 0.54 | 1.00 | 0.60 |
| **ATAC umap** | 0.50 | 0.35 | 0.28 | 0.52 | 0.90 | 0.51 |
| **multigrate umap** | 0.43 | 0.40 | 0.29 | 0.51 | 0.83 | 0.49 |
| **multigrate** | 0.49 | 0.37 | 0.21 | 0.50 | 0.84 | 0.48 |

B

| Method | Bio conservation: Isolated labels | Bio conservation: KMeans NMI | Bio conservation: KMeans ARI | Bio conservation: Silhouette label | Bio conservation: cLISI | Aggregate score: Bio conservation |
|---|---|---|---|---|---|---|
| **MBTA umap** | 0.56 | 0.54 | 0.42 | 0.64 | 0.98 | 0.63 |
| **RNA umap** | 0.57 | 0.55 | 0.40 | 0.63 | 0.99 | 0.63 |
| **MBTA** | 0.52 | 0.52 | 0.37 | 0.52 | 0.98 | 0.58 |
| **totalVI** | 0.53 | 0.40 | 0.29 | 0.54 | 0.99 | 0.55 |
| **protein umap** | 0.50 | 0.38 | 0.30 | 0.57 | 0.95 | 0.54 |
| **totalVI umap** | 0.51 | 0.40 | 0.27 | 0.51 | 0.97 | 0.53 |
| **multigrate** | 0.49 | 0.27 | 0.18 | 0.55 | 0.98 | 0.49 |
| **multigrate umap** | 0.47 | 0.35 | 0.24 | 0.44 | 0.96 | 0.49 |

Figure S8: Quality of embedding generated using various methods under various metrics for the mouse brain (A) and the HSPC (B) datasets.

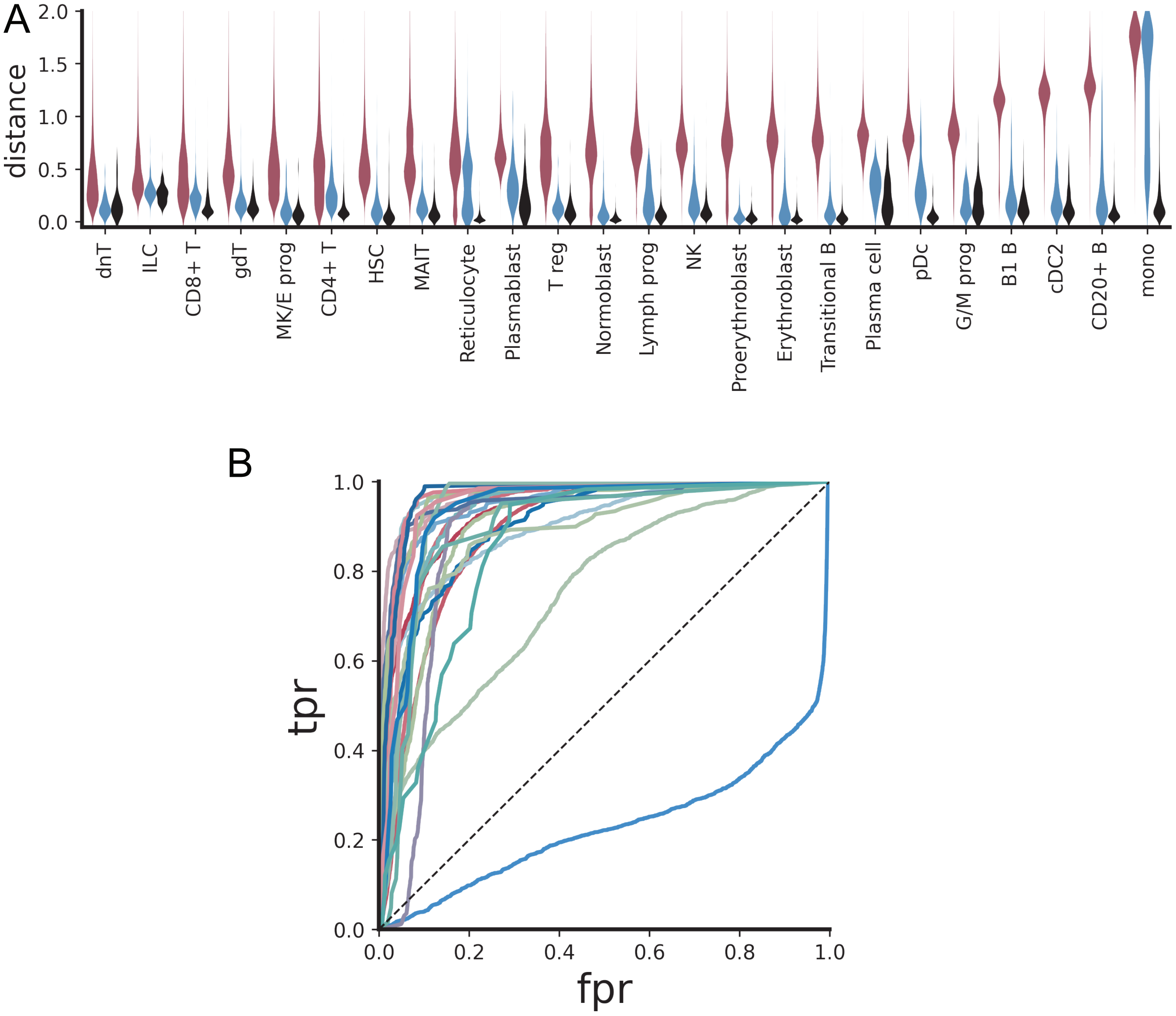


Figure S9: Distribution of distances between each cell and each cluster centroid in the protein space (A), and the receiver operating characteristic curve obtained using these distances to classify the cluster of origin for each cell. Curves are colored by cell type.

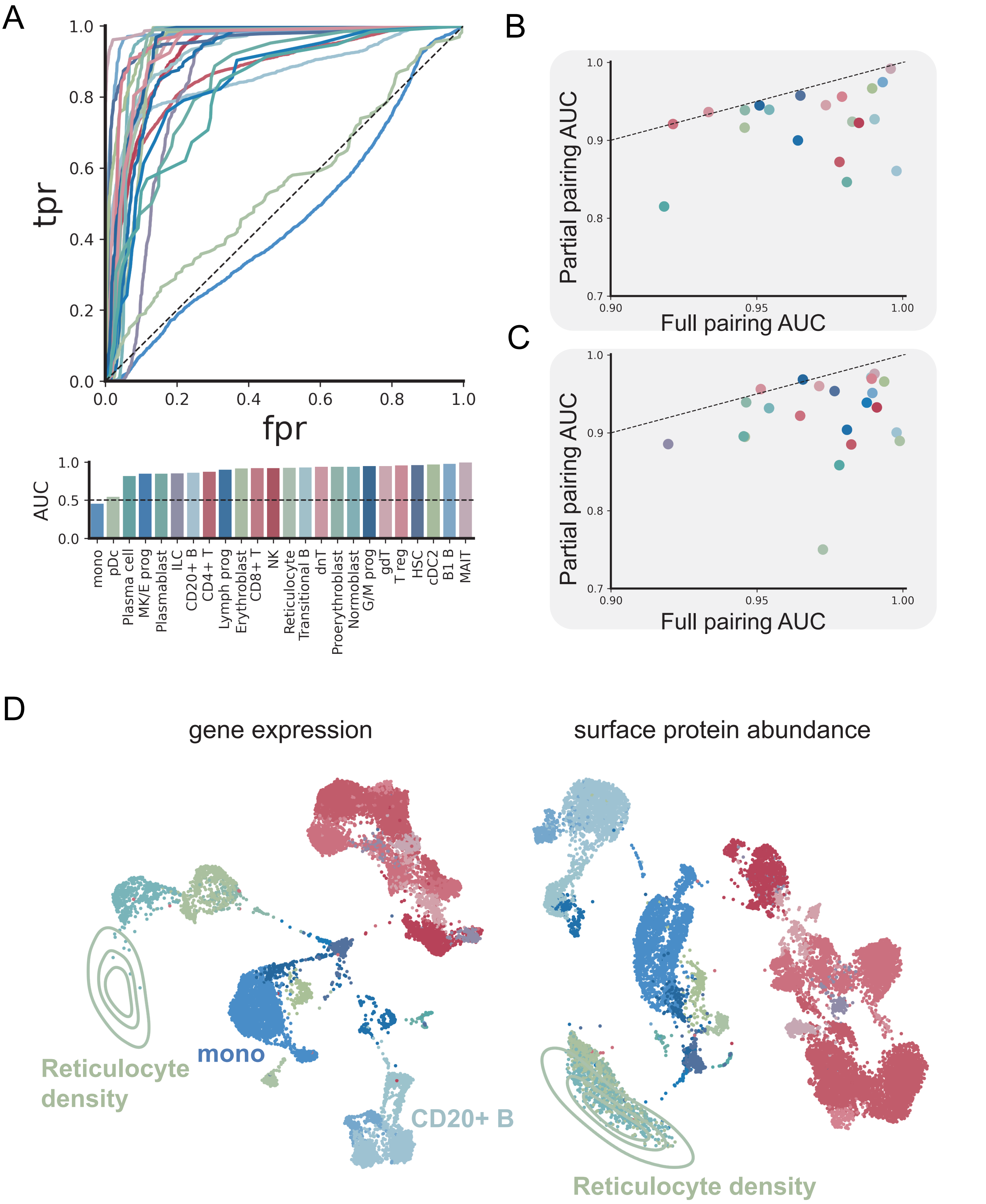


Figure S10: A: Distances between a MBTA translated cell to ground truth cluster centroid are used to predict the originating cell type of each cell and the results are quantified via the receiver operating characteristic curve. B,C: AUCs computed from the partial pairing model and full pairing model from the RNA space (B) and protein space (C). D: UMAP projection of the gene expression (left) and the surface protein abundance (right) data, illustrating that gene expression of cells belonging to the erythroblast lineage is different while their surface protein expression is near identical to each other

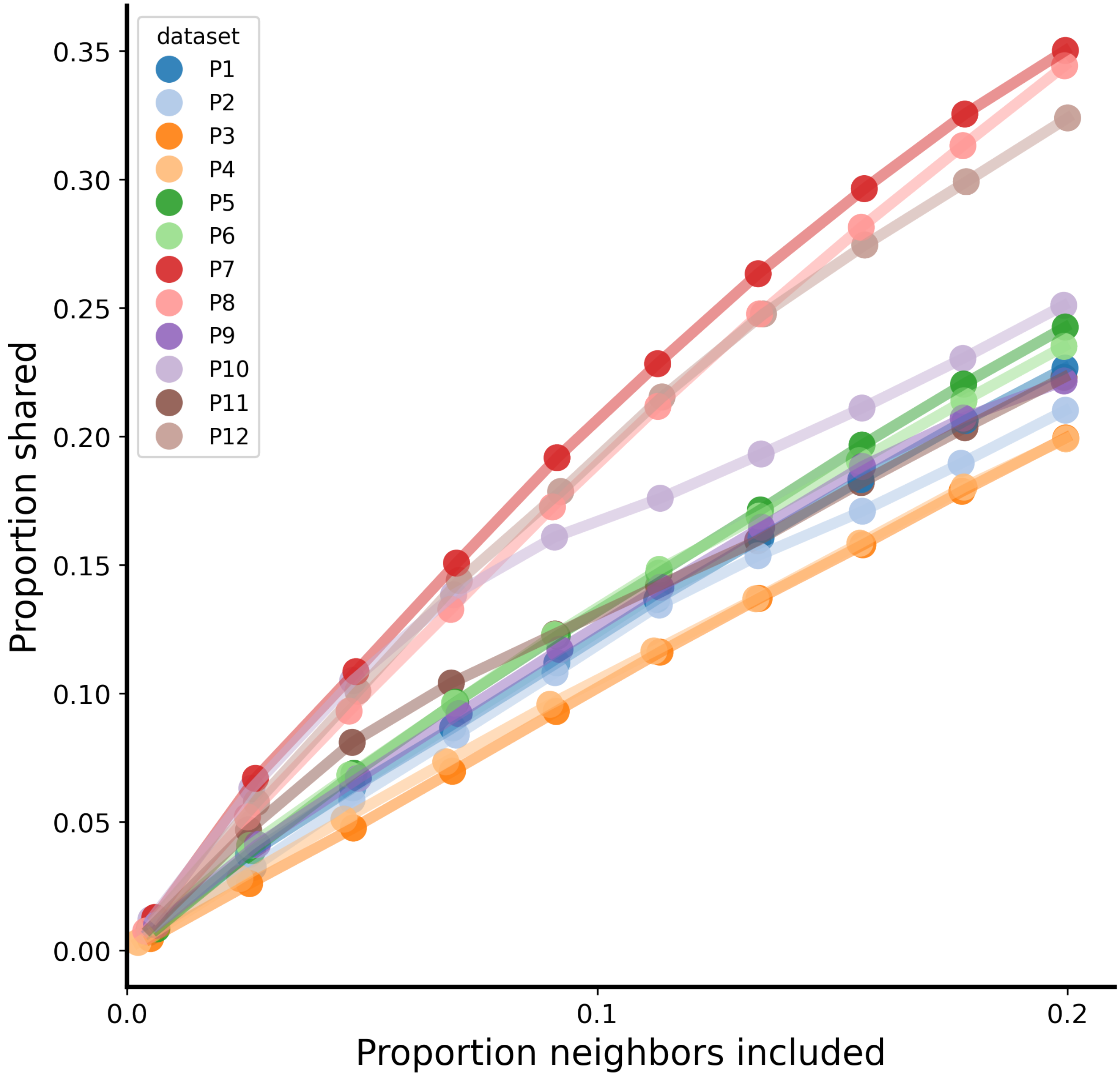


Figure S11: The proportion of neighbors shared between the RNA space and the DNA space, plotted against the proportion of neighbors included of 12 ER+ breast tumor patients [17].

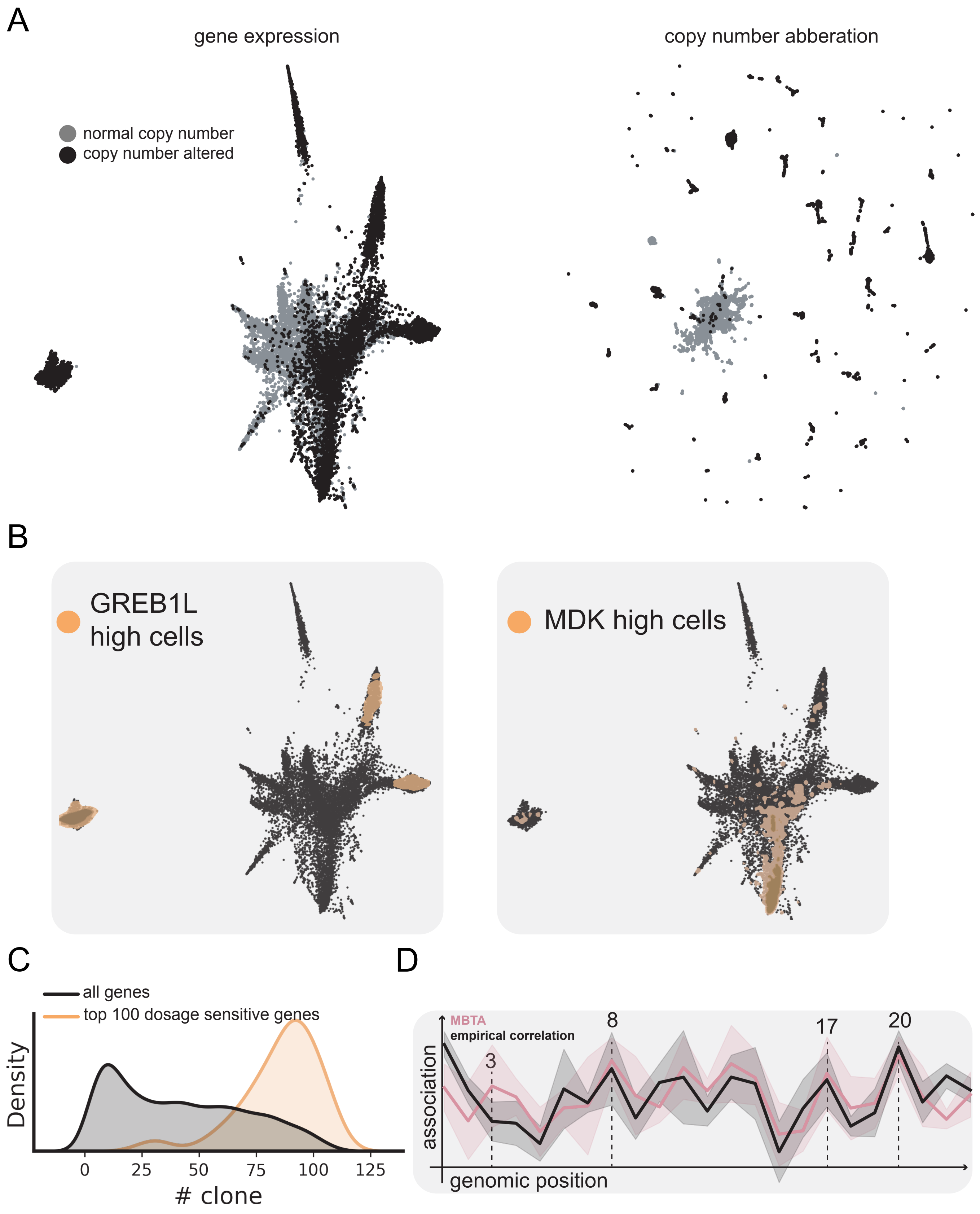


Figure S12: A: UMAP projection of the gene expression (left) and copy number (right) profiles of all twelve patients. B: Cells that highly express exemplary dosage-sensitive genes visualized on the RNA UMAP. C: The distribution of the number of clones from which cells highly express each dosage-sensitive gene. D: Inferred association between chromosomal region and gene expression change by MBTA (pink) and empirical correlation between gene expression and copy number profile (black). Results are averaged over the entire chromosome and the shaded area represents one standard deviation. Example chromosomes with high association with respect to gene expression are labeled.

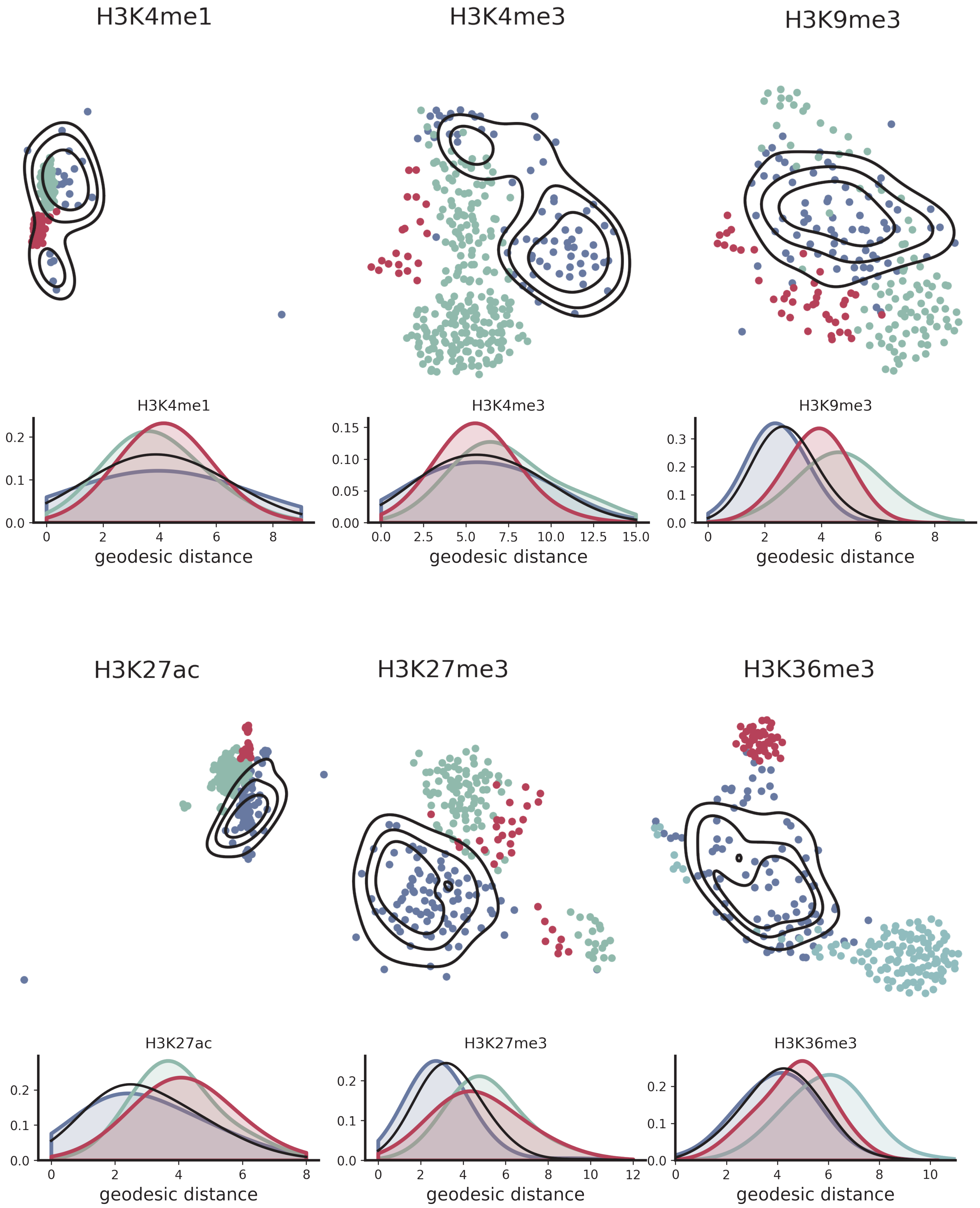


Figure S13: UMAP co-embedding of observed and model-constructed epigenetic features of cells of the 4-cell stage in the mouse embryonic development dataset.

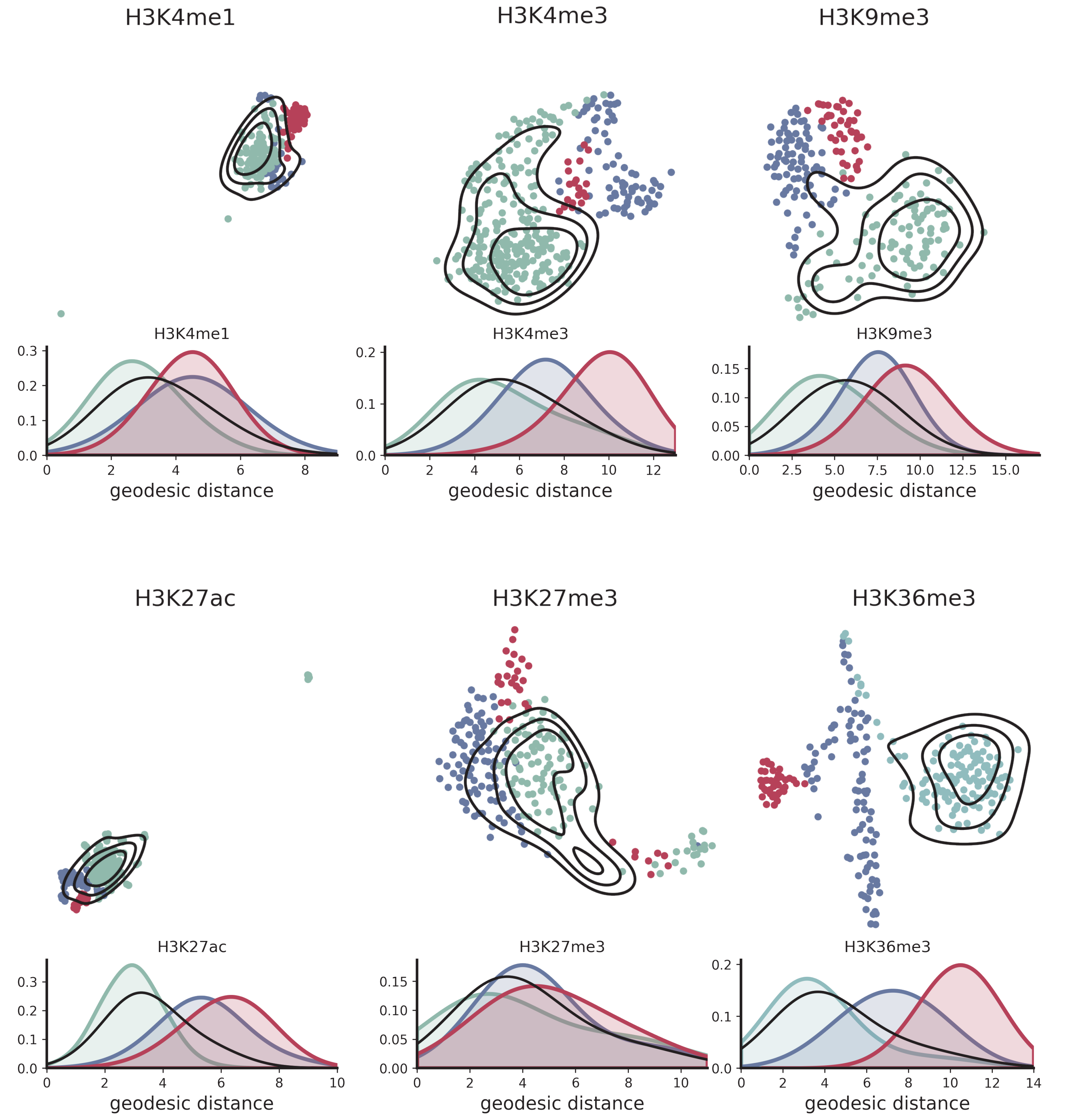


Figure S14: UMAP co-embedding of observed and model-constructed epigenetic features to the E3.0 stage in the mouse embryonic development dataset.

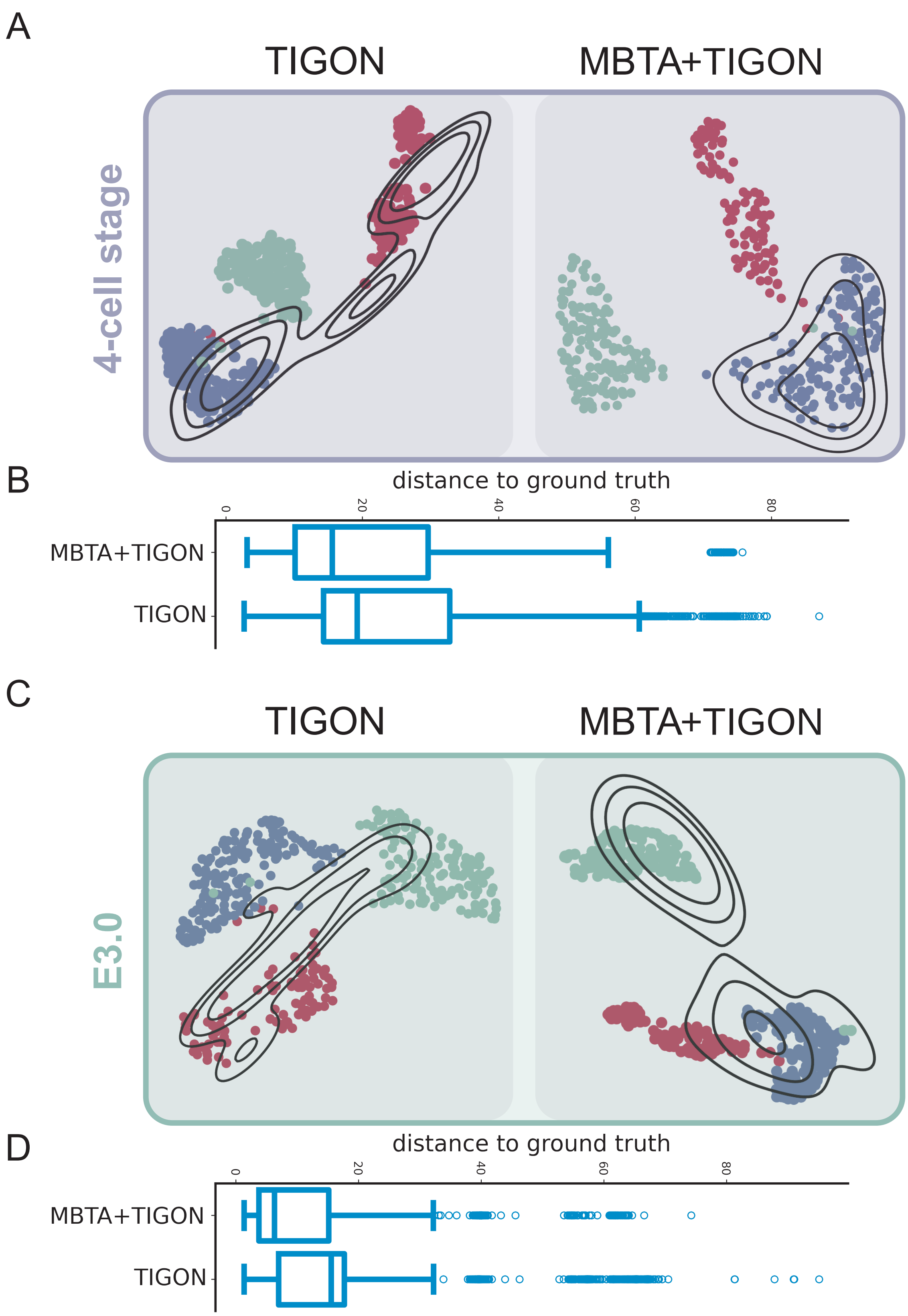


Figure S15: The gene expression of zygotic cells are evolved to the 4-cell stage and the E3.0 stage either directly via TIGON or their H2A.Z profiled are first evolved via TIGON and then translated to gene expression via MBTA. UMAP projection of the zygotic cells evolved using TIGON directly or the "evolve and translate" strategy to the 4-cell stage (A) or the E3.0 stage (C). Distance quantification between the ground truth 4-cell stage cells (B) or the ground truth E3.0 stage cells (D) and the evolved zygotic cells.

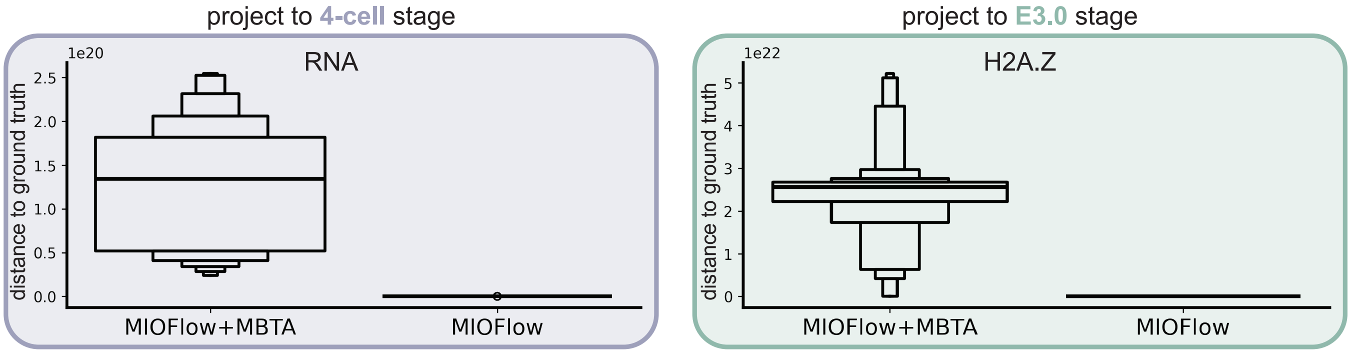


Figure S16: MIOFlow is used to construct unimodal flow for RNA and H2A.Z. The boxplot quantifies the distance between model-generated and target samples using either the "evolve and translate" strategy or MIOFlow directly. Zygotic cells are used as input to construct the RNA profile and the H2A.Z profile of cells of the 4-cell stage and E3.0 stage, respectively.

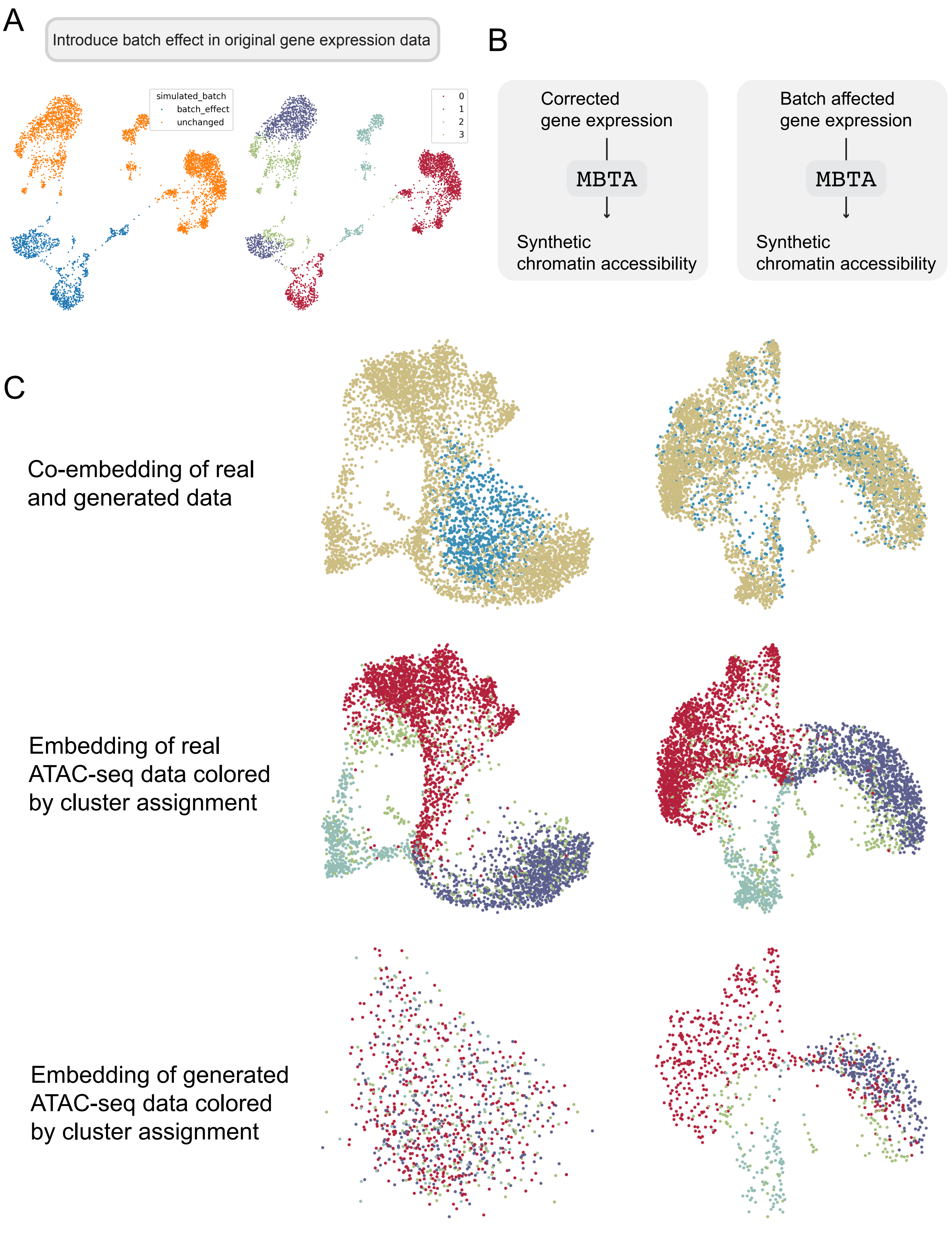


Figure S17: Caption on next page.

Figure S17: A: UMAP visualization of the unchanged and the batch-affected data on the same UMAP plot (left panel); and the same UMAP visualization colored by cell type annotation (right panel). The dataset comes from a co-profiled PBMCs. B: scVI is used to correct for batch effect and the corrected gene expression is used as input to MBTA to generate a synthetic chromatin accessibility profile. As a comparison, batch affected data is also supplied to MBTA to generate an accessibility profile. C: UMAP projection of the synthetic accessibility profiles with (right) and without (left) scVI correction.

| Method | Bio conservation: Isolated labels | Bio conservation: KMeans NMI | Bio conservation: KMeans ARI | Bio conservation: Silhouette label | Bio conservation: cLISI | Aggregate score: Bio conservation |
|---|---|---|---|---|---|---|
| **batch corrected** | 0.59 | 0.39 | 0.40 | 0.58 | 0.87 | 0.56 |
| **batch affected** | 0.48 | 0.01 | 0.00 | 0.47 | 0.41 | 0.28 |

Figure S18: Quantitative evaluation of MBTA-generated ATAC-seq profiles under batch-corrected and batch-affected conditions. Biological conservation metrics for MBTA-generated ATAC-seq profiles evaluated on a PBMC query dataset, comparing a batch-corrected condition (top row) and a batch-affected condition (bottom row, synthetic per-gene offsets introduced).

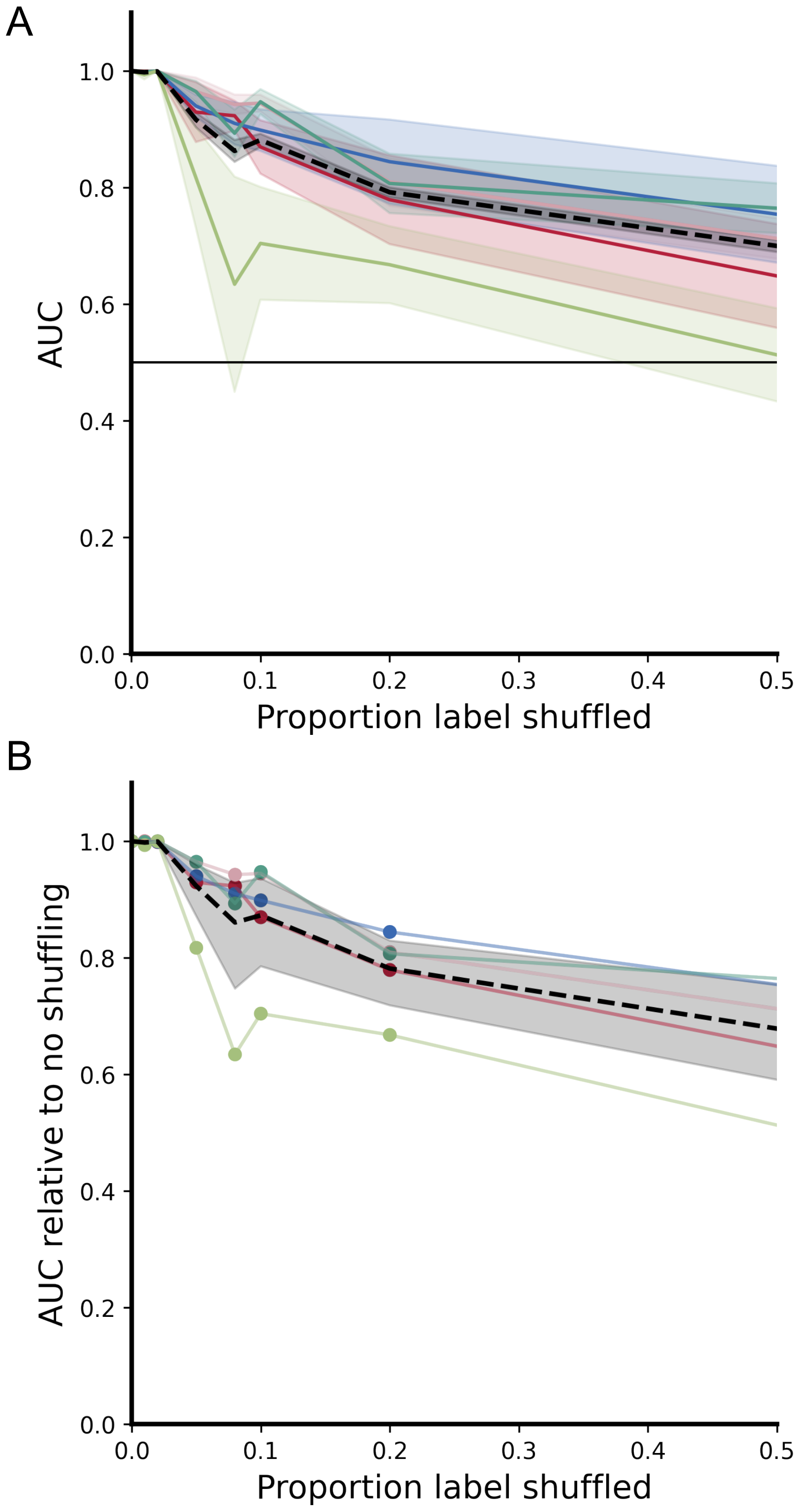


Figure S19: A: Classification accuracy AUC when MBTA model is trained with a proportion of the cell type labels shuffled. Distance between translated gene expression and centroid of each annotated cluster is used for cluster assignment. Color denotes annotated cell type cluster. B: Per cell type AUC relative to when all cell type labels are correct. Black dashed line represents ensmble average. The synthetic cluster dataset is used here.

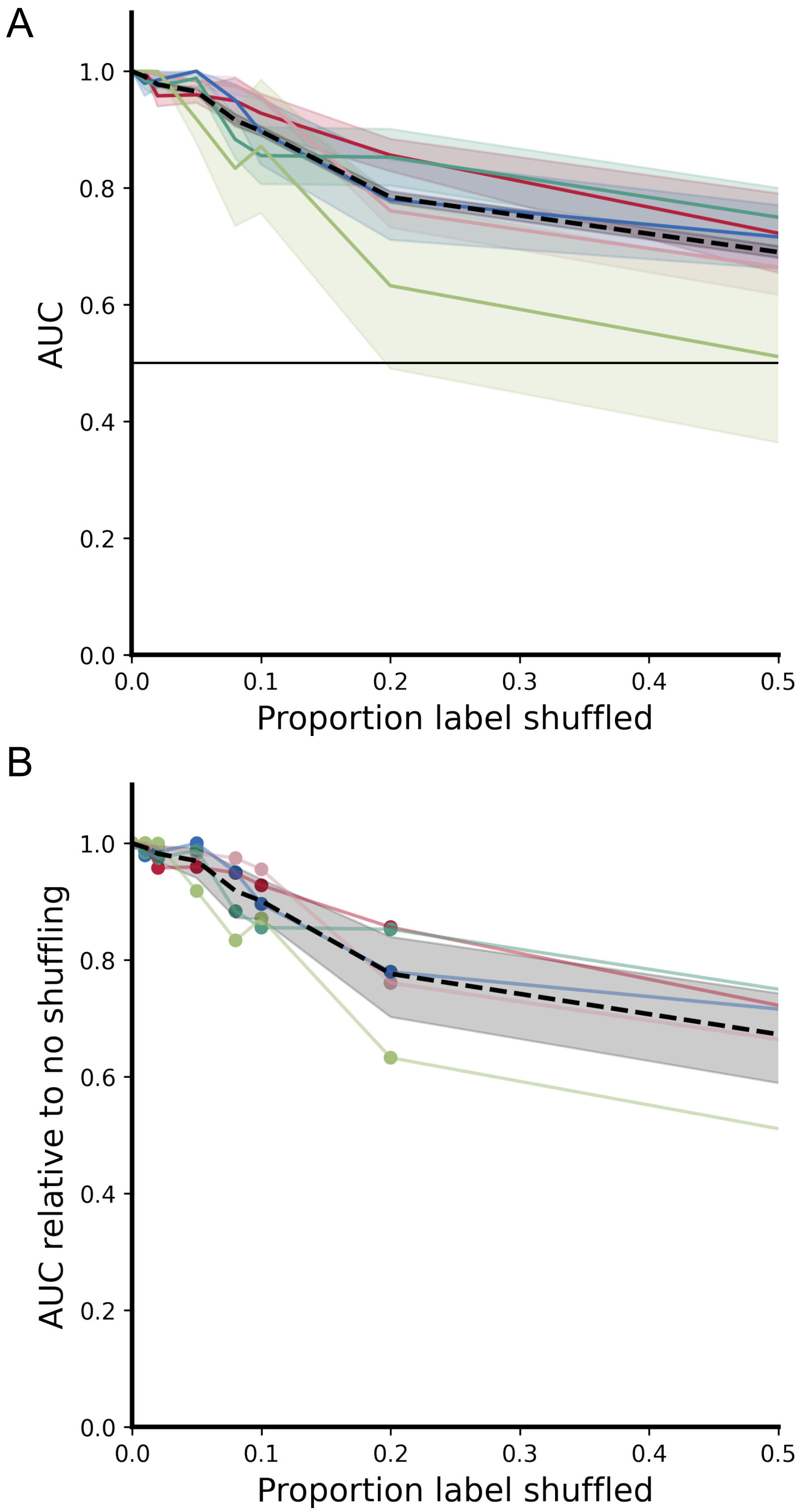


Figure S20: A: Classification accuracy AUC when MBTA model is trained with a proportion of the cell type labels shuffled. Distance between translated chromatin accessibility and centroid of each annotated cluster is used for cluster assignment. Color denotes annotated cell type cluster. B: Per cell type AUC relative to when all cell type labels are correct. Black dashed line represents ensmble average. The synthetic cluster dataset is used here.

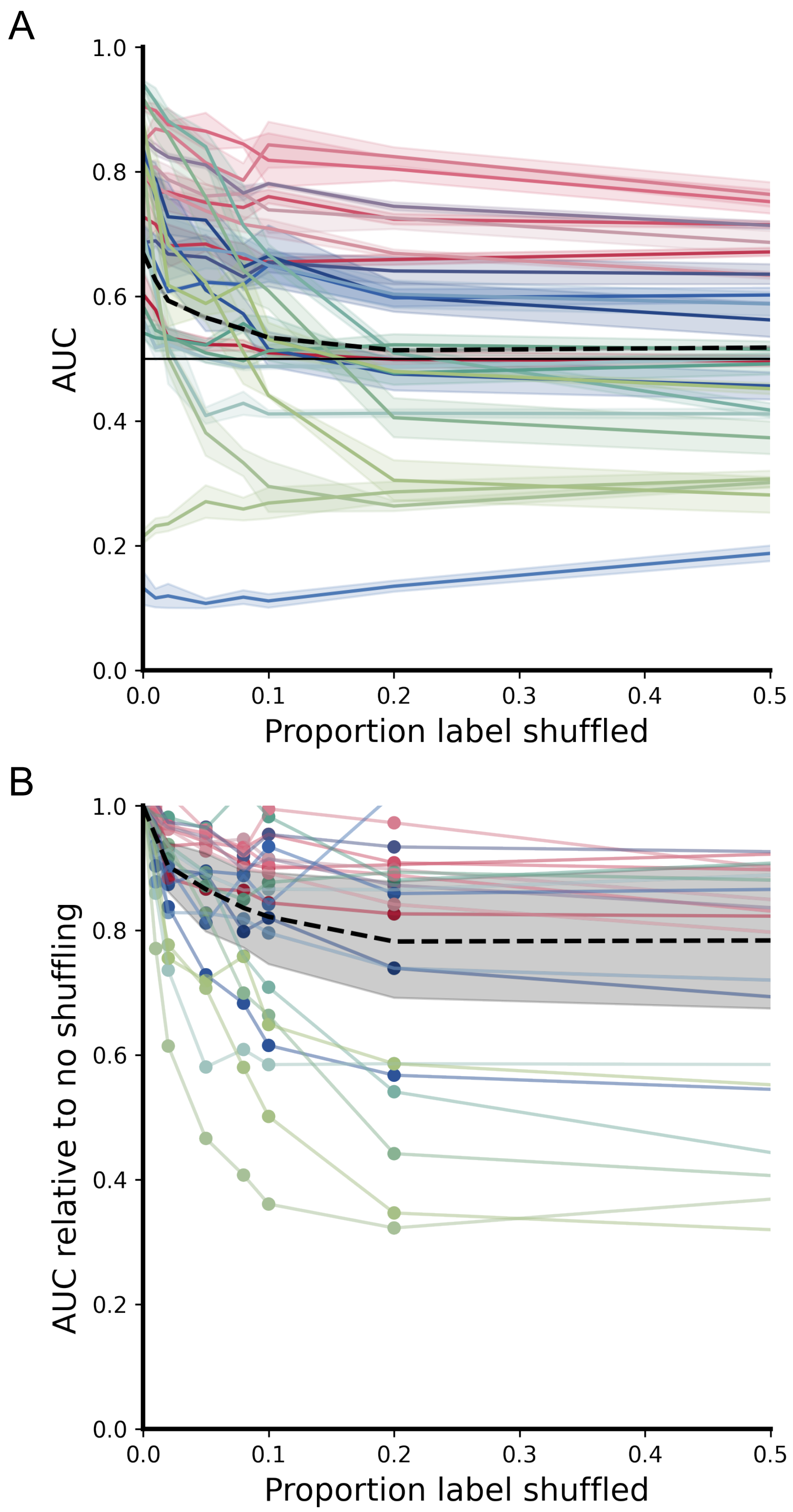


Figure S21: A: Classification accuracy AUC when MBTA model is trained with a proportion of the cell type labels shuffled. Distance between translated gene expression and centroid of each annotated cluster is used for cluster assignment. Color denotes annotated cell type cluster. B: Per cell type AUC relative to when all cell type labels are correct. Black dashed line represents ensmble average. The BMCC dataset is used here.

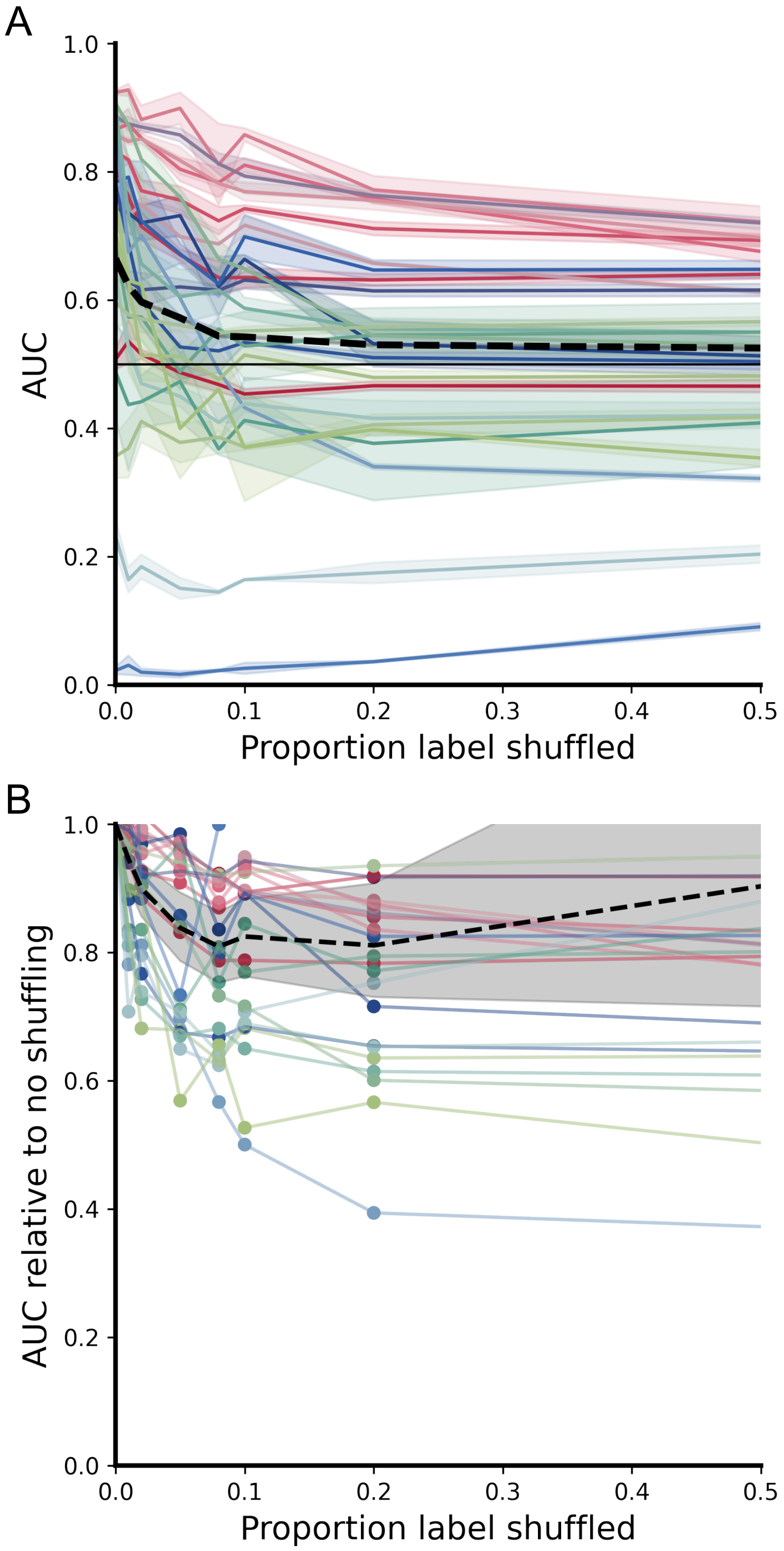


Figure S22: A: Classification accuracy AUC when MBTA model is trained with a proportion of the cell type labels shuffled. Distance between translated protein abundance and centroid of each annotated cluster is used for cluster assignment. Color denotes annotated cell type cluster. B: Per cell type AUC relative to when all cell type labels are correct. Black dashed line represents ensmble average. The BMCC dataset is used here.

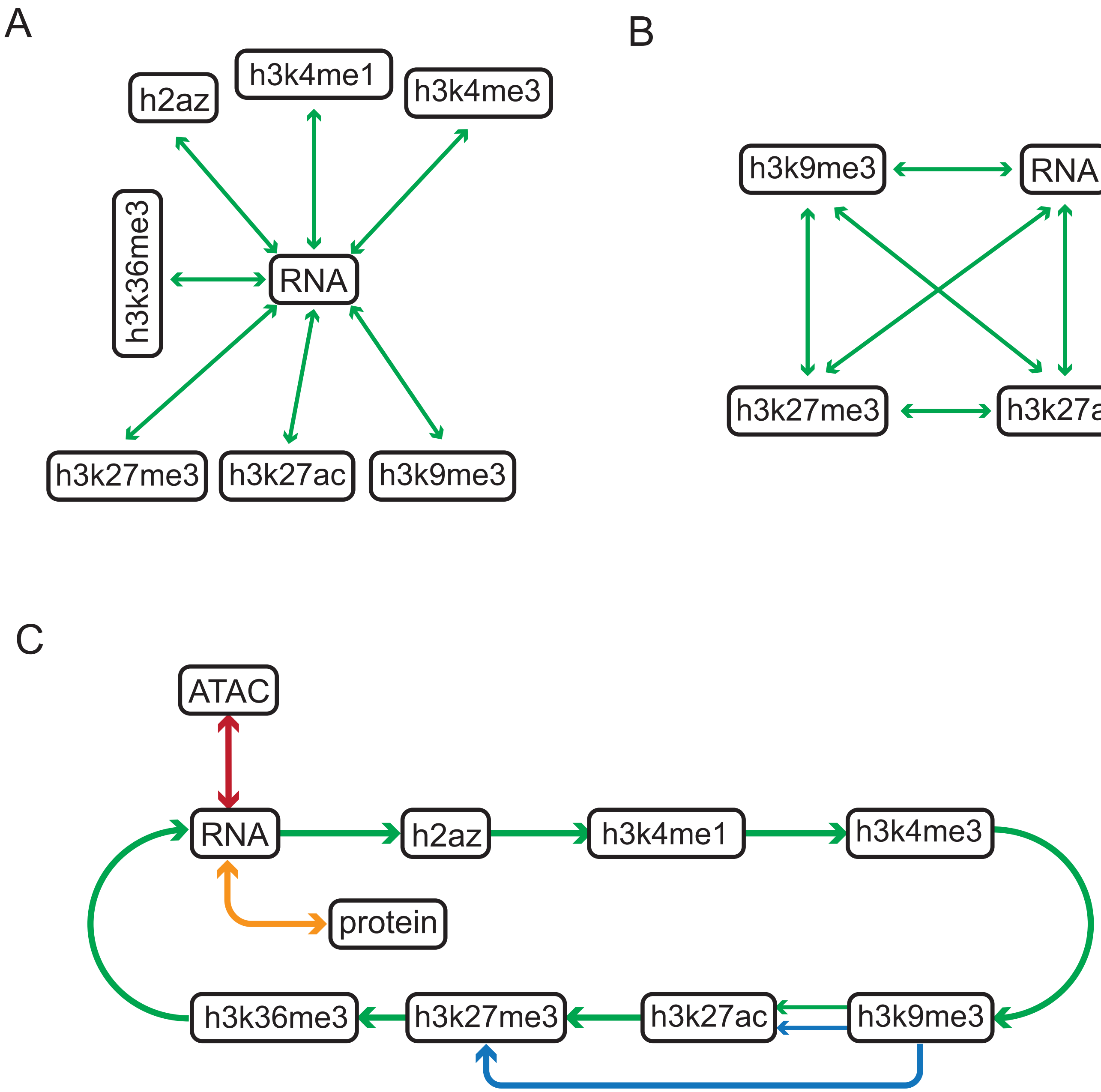


Figure S23: Example station design allowed by MBTA. Each node represents a modality (a trained VAE latent space) and each directed edge represents a trained flow matching component connecting a source modality to a target modality. (A) Hub design. RNA serves as the central hub modality. All other modalities are connected to RNA via direct bidirectional edges where any modality can be translated to any other in at most two steps — first to RNA, then out to the target — at the cost of introducing one intermediate step for non-RNA pairs. (B) All-to-all design. Every modality is connected directly to every other modality via bidirectional edges. Every translation is a single step, minimizing accumulated error, but the number of required components grows quadratically with the number of modalities. (C) Hybrid circular design. Modalities are arranged in a ring with additional direct connections for high-priority pairs (shown in red and orange). Direct edges are reserved for modality pairs that are translated most frequently or for which accuracy is critical; remaining pairs are reached via the ring. This design balances efficiency and accuracy, and illustrates the modular nature of MBTA: new direct connections can be added at any time by training a single new flow model, without retraining any existing components.
